# Massively parallel characterization of adolescent idiopathic scoliosis risk variants

**DOI:** 10.64898/2026.08.05.743047

**Authors:** Darius Ramkhalawan, Justin Koesterich, Fahim Rejanur Tasin, Carlos Cuna, Anat Kreimer, Nadja Makki

**Author notes:** These authors contributed equally to this work (Darius Ramkhalawan, Justin Koesterich, Anat Kreimer, Nadja Makki).

## Abstract

Adolescent idiopathic scoliosis (AIS) is a common pediatric musculoskeletal disorder characterized by lateral spinal curvature, often leading to chronic pain and deformity. While a significant genetic component to AIS is recognized, the functional impact of most associated genetic variants, particularly those in non-coding regions, remains largely unknown.

Using massively parallel reporter assays, we characterize 1,664 variant positions in linkage disequilibrium with 26 AIS lead variants identified by genome-wide association studies (GWAS) in chondrocytes, a major cell type implicated in AIS pathogenesis. Using a library of 7,173 candidate regulatory sequences we compare the 1,664 reference alleles against 4,708 alternate alleles in two human chondrocyte cell lines (TC28a2 and SW1353). Our analysis identifies 92 variants that exhibit significant differential regulatory activity between their reference and alternate alleles, 79 of which are predicted to disrupt transcription factor binding sites, often correlating with their observed regulatory effect.

Notably, we validate rs9496392, a single-nucleotide variant near the *ADGRG6* locus, which shows consistent differential regulatory activity in both cell lines. ADGRG6 is a key regulator of cartilage homeostasis, and its cartilage-specific knockout in mice results in a scoliosis-like phenotype. The AIS risk allele of rs9496392 (T) is predicted to strongly disrupt several TFBSs, including SP1.

This study provides a foundational catalog of functional AIS-associated regulatory variants active in chondrocytes, offering crucial insights into the perturbed gene regulatory networks in AIS. These findings lay the groundwork for identifying biomarkers and potential therapeutic targets for this complex childhood disease.

## INTRODUCTION

Adolescent idiopathic scoliosis (AIS), a lateral spinal curvature spontaneously developing during puberty in otherwise healthy children, is the most common pediatric musculoskeletal disorder globally (E J Rogala et al. 1978; Weinstein 2019; C. Wise et al. 2008). The progression of AIS can severely impact a patient’s life, leading to chronic pain, physical deformity, spinal osteoarthritis, and impaired lung and heart function (Hoelen et al. 2023). Current treatment options are limited to bracing or corrective surgery, and the absence of pre-symptomatic diagnostic capabilities underscores the need for a deeper mechanistic understanding of the disease.

Genetic studies have established a significant genetic component to AIS (Ogura et al. 2015; Karner et al. 2015; Liu et al. 2021; Gray et al. 2021; Yonezawa et al. 2020; Ushiki et al. 2024; Blecher et al. 2017; Patten et al. 2015; Buchan, Alvarado, et al. 2014; Haller et al. 2016; Grauers et al. 2013; Andersen et al. 2007; Yang et al. 2012; Tang et al. 2012; Tuncay et al. 2025), with numerous susceptibility loci identified through genome-wide association studies (GWAS) (Ogura et al. 2015; C. A. Wise et al. 2020; Khanshour et al. 2018; TSRHC Scoliosis Clinical Group et al. 2015; Londono et al. 2014; Miyake et al. 2013; Sharma et al. 2011; Sudmant et al. 2015; Kou et al. 2019, 2013; Zhu et al. 2015; Ogura et al. 2017; Wu et al. 2019; H. Yu et al. 2024; Takahashi et al. 2011; Fan et al. 2012; Japan Scoliosis Clinical Research Group (JSCRG) et al. 2018). However, the vast number of potential causal variants, especially those in linkage disequilibrium (LD) with GWAS hits (exceeding one thousand), presents a major hurdle. Most of these variants are located in non-coding regions, suggesting their involvement in regulating gene activity. A few studies have begun to identify AIS-associated variants with altered regulatory function through individual enhancer assays (TSRHC Scoliosis Clinical Group et al. 2015; Yonezawa et al. 2020). It was also recently demonstrated that knockout of two enhancer elements harboring AIS-associated variants at the *PAX1* locus, results in a kinked tail phenotype in mice (Ushiki et al. 2024). However, the comprehensive identification of functional variants at AIS loci remains an outstanding challenge.

Compelling evidence supports the involvement of abnormal cartilage biogenesis and development in AIS onset. Recent studies suggest that estrogen signaling may disrupt extracellular matrix homeostasis in growth plate cartilage (H. Yu et al. 2024), and cartilage-specific deletion of the AIS susceptibility genes *ADGRG6* in mice has been shown to induce a scoliotic curve during adolescence (Karner et al. 2015; Liu et al. 2021). Despite these insights from human genetic studies and animal models, a critical gap exists in understanding how the non-coding variants identified in patients contribute mechanistically to AIS pathogenesis.

In this study, we carried out the first massively parallel reporter assay (MPRA) to comprehensively assess the regulatory activity of AIS-associated variants. This innovative, high-throughput approach allowed for an unbiased examination of all AIS GWAS variants (Tang et al. 2012) and those in LD in a single quantitative experiment. Given the central role of chondrocytes in AIS pathogenesis, MPRAs were performed in two human chondrocyte cell lines. This work provides a highly confident list of functional variants and their regulatory effects. The functional variants identified in this study will serve as a foundational resource for elucidating altered gene regulatory networks in AIS patients and will significantly advance the future investigations of this common childhood disease. Importantly, this work lays the groundwork for the identification of AIS biomarkers and potential drug targets.

## RESULTS

### MPRA variant library design and selection of cell lines

We designed our MPRA library to include all 26 lead AIS GWAS SNPs (C. A. Wise et al. 2020), as well as variants in LD (r2 ≥ 0.7) in European, Han Chinese, and ethnic Japanese populations resulting in 1,664 unique variants (1,522 SNPs, 142 indels) (Supplemental Table 1). Each candidate regulatory sequence (CRS) was generated as a 200 base pair sequence with the variant of interest at base pair 101. For each variant we included all possible alleles (4 per SNP, 2 per indel) resulting in a total of 6,372 test CRSs. Given the central role of cartilage in AIS pathogenesis (Karner et al. 2015; Liu et al. 2021), we sought to conduct our MPRA in two commonly used human chondrocyte cell lines: TC28a2 and SW1353. For positive controls, we identified regions of significant H3K27ac enrichment and filtered peaks according to RNA-seq expression in both cell lines, where only sequences within 1 Mb of a gene with expression of TPM > 1 were retained, resulting in 601 positive control sequences. As negative controls, we included 200 scrambled sequences from randomly selected reference candidate regulatory sequences. This resulted in a total library size of 7,173 CRS, including those with AIS variants to be tested for regulatory activity, as well as positive and negative controls (Figure 1).

**Figure 1:**
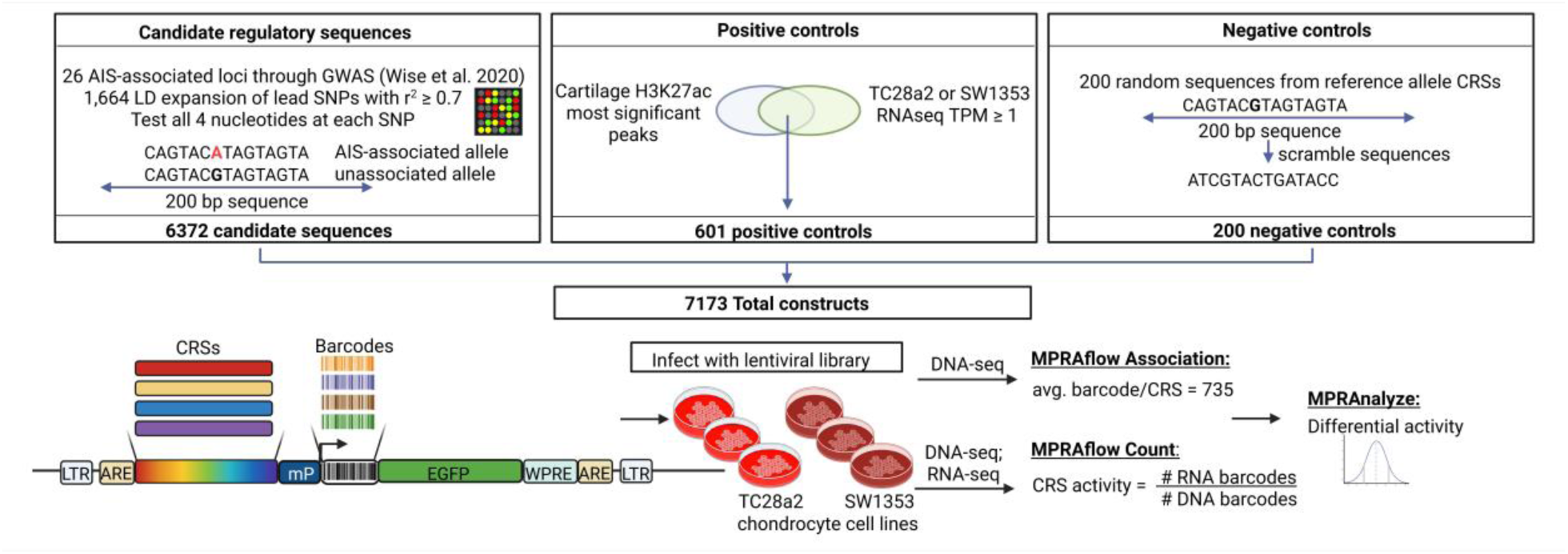
Overview of MPRA design generation. This schematic shows the selection process of choosing the candidate regulatory sequences to be investigated in the assay (top left). selection of positive controls (top middle) and negative controls (top right). Bottom shows an overview of the MPRA steps including the segment of DNA that is integrated into the cell genome (bottom left), transduction into chondrocyte cell lines (bottom middle) and the downstream sequencing and processing of the MPRA (bottom right).

### MPRA association, count, and significant variants pairs

A minimal promoter, random barcode sequences, and vector overhangs were added to CRS oligonucleotides by sequential rounds of PCR and cloned into a lentiMPRA reporter vector. To associate CRSs with their respective barcodes, we sequenced the CRS-barcode fragment and utilized the MPRAflow pipeline to associate the randomly combined transcribable barcodes with the upstream CRS (Gordon et al. 2020). We identify over 7.3 million barcodes confidently assigned to our CRSs at an average of 735 barcodes per CRS.

Next, we infected each cell line with the lentivirus library in triplicate, then simultaneously isolated DNA and RNA for sequencing of DNA and RNA barcodes. Active CRSs drive transcription of the downstream barcode sequences; the transcribed RNA barcodes are then normalized to the integrated DNA barcodes to quantify the gene regulatory activity of the CRS. We analyzed the DNA and RNA obtained from the infected chondrocytes and matched the counts for the barcodes with those that were associated with the CRSs. We were able to match counts for 66.27% of the confidently associated barcodes in the TC28a2 cell line. We observed an average r correlation value of 85% for the DNA barcode counts between replicates and 80.33% for RNA barcode counts between replicates (Supplemental Figure 1). For the SW1353 cell line, we confidently associated 61.98% of barcodes and observed an average r correlation value of 87.33% for the DNA barcode counts between replicates and 77% for RNA barcode counts between replicates (Supplemental Figure 2). When aggregating replicate counts to the CRS level, we observe approximately 98% DNA, 96% RNA, and 92% RNA/DNA ratio r correlation values between replicates in both cell lines (Supplemental Figures 3 and 4).

After quality control steps were performed, we retained 6,266 CRSs for TC28a2 and 6,265 CRSs for SW1353, plus 791 positive and negative controls, for further statistical analysis. With a total of 4,878,602 and 4,563,151 confidently counted barcodes, for TC28a2 and SW1353 respectively. Next, we utilized the MPRAnalyze program (Ashuach et al. 2019) to generate estimated transcription rates from the sequenced RNA and DNA. These transcription rates, referred to as alpha values, are first compared to the negative controls to identify sequences that are significantly active in the cell line. We identified a total of 1,566 CRSs in TC28a2; 911 in SW1353; and 2,008 combined across both cell lines (659, 499, and 886 unique loci respectively) that had a significantly active transcription rate compared to the negative controls (*p*-value ≤ 0.05; Figure 2 A, B; Supplemental Tables 2 and 3). This represents the first functional high-throughput identification of active enhancer elements in chondrocytes and will serve as a new catalog of active enhancers in TC28a2 and SW1353 cells.

**Figure 2:**
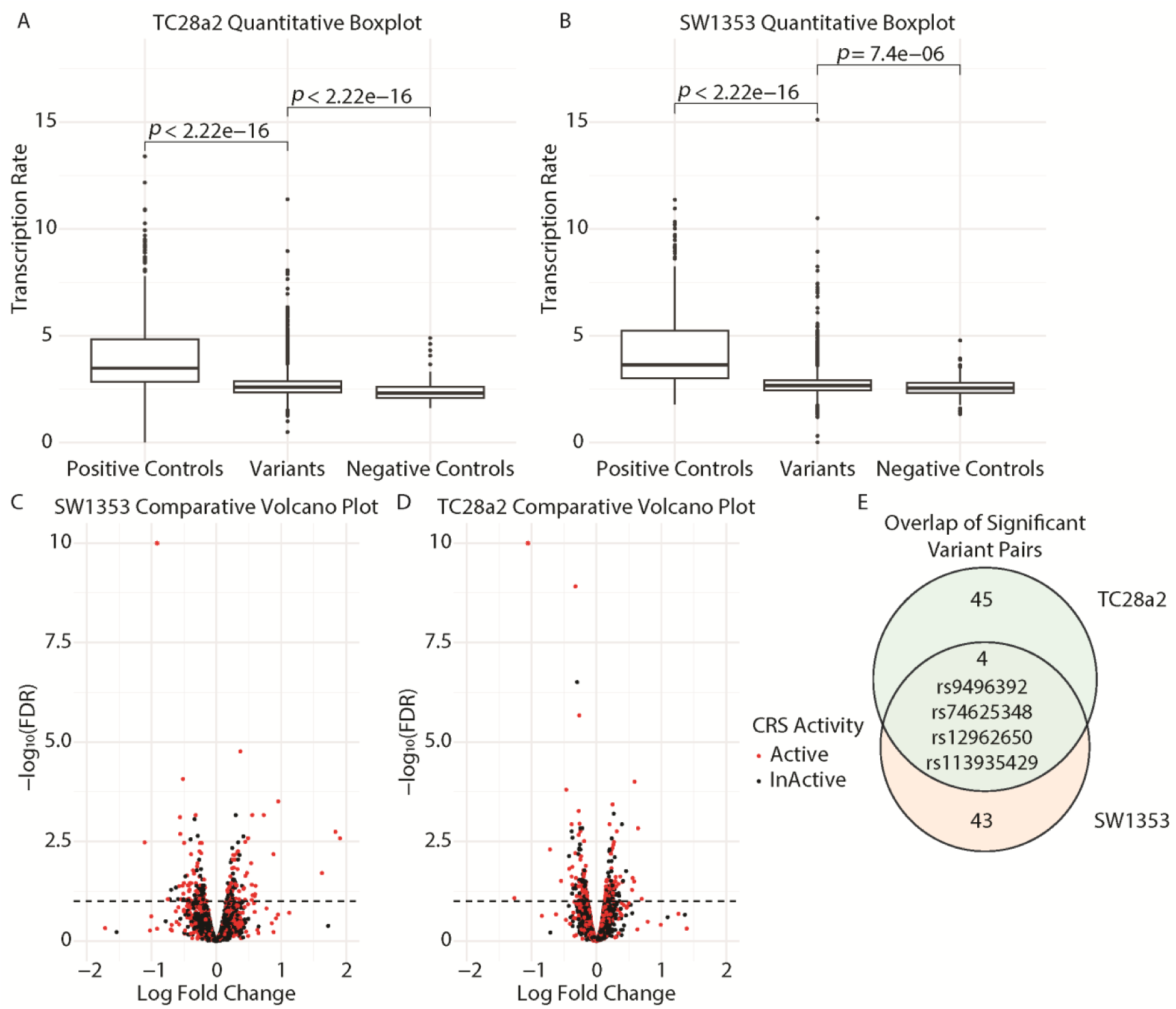
Quantitative and comparative MPRA results. A, B) Boxplots showcasing the Quantitative analysis performed on the RNA/DNA transcription rate for all CRSs, separated into positive controls, negative controls, and combined reference and alternate allele (Variants). *p*-values at the top of the boxplots were calculated using Wilcoxon rank-sum test comparing the median transcription rate between the 2 conditions listed under the bracket. A separate *p*-value calculation is done by MPRAnalyze to determine individual CRSs that are active compared to the negative controls (Methods). C, D) Volcano plots showing the Comparative analysis performed by MPRAnalyze on the reference alternate variant pairs. Each point is a single Reference/Alternate variant pair, positioned based on the comparative analysis and colored based on the quantitative analysis. The x-axis denotes the natural log fold change values of the CRS; positive LFC indicates a higher transcription rate in the reference allele compared to the alternate allele. The y-axis denotes the -log_10_ of the FDR value that is associated with the variant pair’s log fold change. A variant pair is called as significantly disruptive if the FDR value is less than 0.10, denoted by the dashed line. Additionally, we call a final significantly disruptive variant pair list as those that are both significantly disruptive, above the dashed line, as well as either the reference or alternate allele is significantly active compared to the negative controls, denoted as being red. E) Overlap of the final significant observed alternate allele variant pairs identified in the 2 cell lines.

MPRAnalyze then groups the CRSs by reference-alternate variant pairs, which are analyzed to identify variant pairs that contain a significant difference in transcription between the two alleles. As our library contains 1,664 unique variants, comparing each SNP reference allele to its three alternate alleles (1,522 SNPs × 3 = 4,566) plus 142 indel pairs results in 4,708 total testable variant pairs. After removing incomplete variant pairs and variant pairs with only zero or missing counts, we investigated the allelic effect of 4,468 variant pairs in both cell lines, representing 1,579 of the starting 1,664 unique variants.

Of the 4,468 retained total variants pairs, we identify 208 total variant pairs in the TC28a2 cell line, along with 192 total variant pairs in the SW1353 cell line, that have significant change in transcription rates between alleles (FDR ≤ 0.1; Figure 2 C, D; Supplemental Tables 4 and 5). These variant pairs included either the GWAS/HapMap alternate variant or a synthetic alternate variant. To identify functional regulatory variants underlying the observed genetic association in the AIS GWAS, we focused our downstream analysis on those variant pairs that included the GWAS/HapMap (referred to as observed) alternate allele. This resulted in 1,579 retained observed alternate allele variant pairs, of which 82 observed alternate variant pairs in the TC28a2 cell line, along with 73 observed alternate variant pairs in the SW1353 cell line, that have significant change in transcription rates between alleles (FDR ≤ 0.1; Supplemental Tables 4 and 5).

We then overlap the two analyses to generate a set of 92 highly confident variant pairs (49 in TC28a2, 47 in SW1353) that have both a significant change in transcription rates between alleles and at least 1 allele significantly active compared to negative controls (Figure 2; Supplemental Tables 6 and 7). Interestingly, 4 variant pairs (rs113935429 (T>-), rs9496392 (T>G), rs74625348 (G>C), rs12962650 (T>G)) were significant in both cell lines (Figure 2E). 49 additional variant pairs (55.7%) are found to have the same direction of effect on transcription. 21 of these 49 variant pairs are found with at least 1 allele active in both cell lines and perhaps being called as differentially non-significant in the 2nd cell line is due to insufficient statistical power, rather than biological effects (Supplemental Tables 6 and 7). These 92 regulatory variants are spread across 18 of the 26 AIS GWAS loci and contain 3 of the lead SNPs themselves (Supplemental Figure 5).

### Annotation of significant variant pairs

As enhancers are typically bound by transcription factors (TFs), we next investigated if these variants are predicted to disrupt transcription factor binding sites (TFBS), which would improve our understanding of how specific variants disrupt transcription. Here we leverage the motifbreakR program (Coetzee et al. 2015) to predict differential binding affinity of TFBSs overlapping the variant position. We found that 79 of the 92 variants (85.9%) are predicted to disrupt the binding site of at least 1 TF that is active in either cell line (Supplemental Tables 6 and 7). Of these 79 variant pairs, 59 have a predicted TF disruption in the same direction of effect as the variant pairs’ MPRA log fold change value. This suggests that the variants disruption to transcriptional regulation is likely due to their effect on transcription factor binding. Interestingly, one of the variant pairs (rs200323221) (Figure 3A) is predicted to disrupt the binding of the transcription factor EGR1 (Figure 3B, C). This variant is shown to have a similar ratio of disruption to both transcription in the MPRA as well as the TFBS predicted binding (Figure 3B). EGR1 has previously been associated with AIS and the disruption of its binding to this potential regulatory region may shed new light on affected transcriptional networks downstream of this TF (Yonezawa et al. 2020).

**Figure 3:**
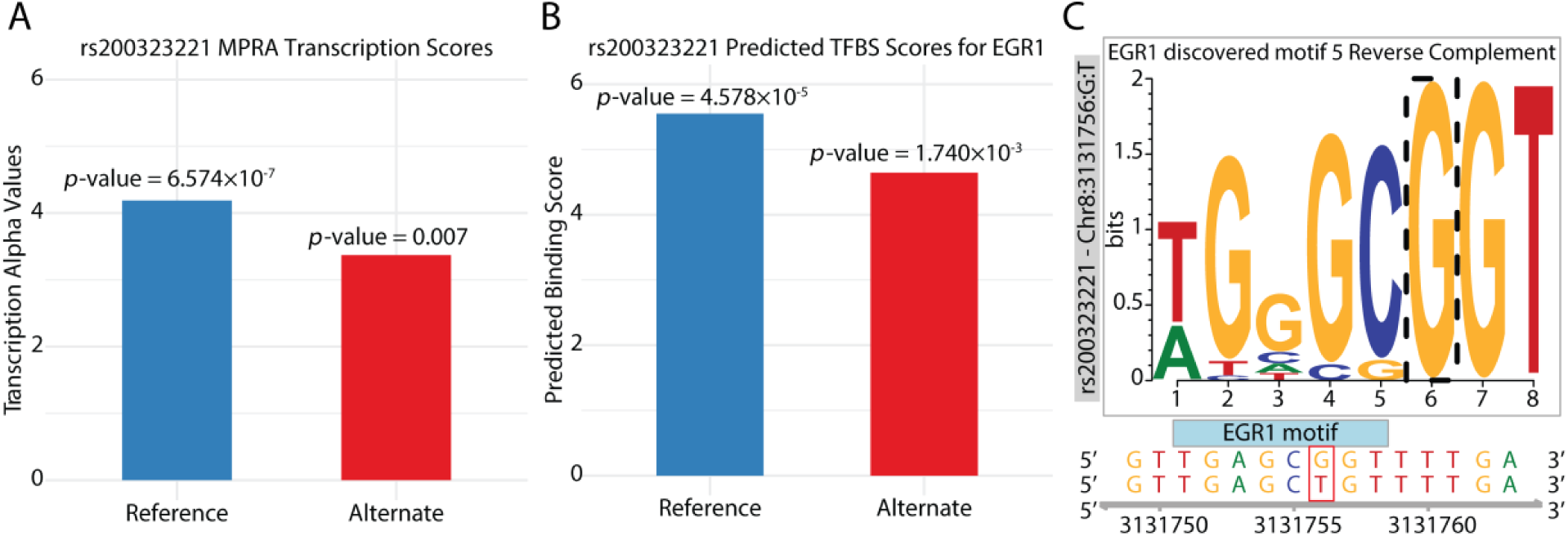
Predicted transcription factor binding site disruptions highlighting predicted TFBS disruptions due to the variant rs200323221. A) MPRA transcription rate (alpha) value for rs200323221 (G>T) in SW1353 cell line. The listed *p*-value is an empirical *p*-value reported by MPRAnalyze for significant activity compared to the negative controls in the SW1353 cell line. B) Predicted binding score for the EGR1 motif with the given allele. The listed *p*-values are calculated based on the likelihood of the sequence similarity score of the allele specific sequence to the TFBS compared to the sequence similarity score using the null background allelic distribution. C) LOGO representation of the predicted motif region with the mutated position (dashed box), reference sequence (below the graph, top row), and the alternate sequence (bottom row), with the reference and alternate alleles highlighted in the red shaded rectangle. The y-axis represents the weight of presence each allele has at that position observed in the ENCODE motif database.

Next, we identified genes closest to the 92 significant variants pairs and performed GO (G. Yu et al. 2012) enrichment analysis. We identify 22 unique proximal genes (Supplemental Tables 6 and 7) which were enriched for skeletal system and connective tissue development, cartilage condensation, regulation of cell differentiation, and cell fate specification (Supplemental Figure 6). When examining expression levels of the 22 genes, we found that 9 are expressed in AIS patient cartilage (TPM ≥ 1; Supplemental Figure 7) (Makki et al. 2021). Next we examined if any of the genes closest to the 92 variants (Supplemental Tables 6 and 7) have previously been linked to AIS through functional studies (Ogura et al. 2015; Karner et al. 2015; Liu et al. 2021; Yonezawa et al. 2020; Buchan, Alvarado, et al. 2014; TSRHC Scoliosis Clinical Group et al. 2015; Ogura et al. 2017; H. Yu et al. 2024; W. Wang et al. 2023; Becker-Heck et al. 2011; Smits and Lefebvre 2003; Y. Wang et al. 2022; Barat-Houari et al. 2016; Buchan, Gray, et al. 2014; Bachmann-Gagescu et al. 2011; Jaffe et al. 2016; de Azevedo et al. 2022; Henry et al. 2012; Van Gennip et al. 2018; Haller et al. 2018; Gray et al. 2014; Rebello et al. 2023; Hoornaert et al. 2010; Hayes et al. 2014; Su et al. 2021; Xu et al. 2025; Grimes et al. 2016; Decourtye et al. 2022; Bieder et al. 2023) (Supplemental Table 8). We identified 40 significant variants in proximity to seven genes previously linked to AIS through functional studies, namely *ADGRG6*, *BNC2*, *FTO*, *MIR4300HG*, *PAX1*, *SOX6*, and *UNCX*. These findings give us potential insights into the mechanisms of both the variant disruption and the downstream involvement in disease pathology.

### Validation and characterization of rs9496392

Among the 92 significant variant pairs identified in our MPRA (Supplemental Table 6 and 7) was a previously reported regulatory variant, rs169311, located within the PEC7 enhancer proximal to *PAX1*. The risk allele (A) was previously shown to abolish enhancer activity in zebrafish (TSRHC Scoliosis Clinical Group et al. 2015), consistent with our MPRA results. We then further validated the regulatory effects of five additional MPRA-identified SNPs using luciferase reporter assays in the TC28a2 cell line (Figure 4A and B, Supplemental Figures 8 and 9). SNPs were selected based on their proximity to genes with roles in chondrogenesis and cartilage homeostasis (*ADGRG6*, *FTO*, *WNT9A*) (Tuncay et al. 2025). Differential luciferase activity of each allele generally coincided with the observed MPRA activity (Figure 4A, 4B, Supplemental Figure 8, Supplemental Figure 9). Some constructs (rs1040525, rs113935429) showed differential luciferase activity in the opposite direction from the MPRA signal, which could potentially be due to the differing context dependencies that are inherent to the two assays(Klein et al. 2020, 2019; Inoue et al. 2017). One of the validated single-nucleotide variants, rs9496392 (T>G), is located at the *ADGRG6* locus (Figure 4A, B). ADGRG6 is a key regulator of cartilage homeostasis and helps maintain proper alignment of the vertebral column during postnatal development (Karner et al. 2015). *ADGRG6* is expressed at high levels in human spinal cartilage, mouse chondrocytes, and mouse intervertebral discs (Makki et al. 2021) (Figure 4C). Additionally, *ADGRG6* is significantly downregulated in AIS patient cartilage tissue compared to control samples (Ramkhalawan et al. 2026).

**Figure 4:**
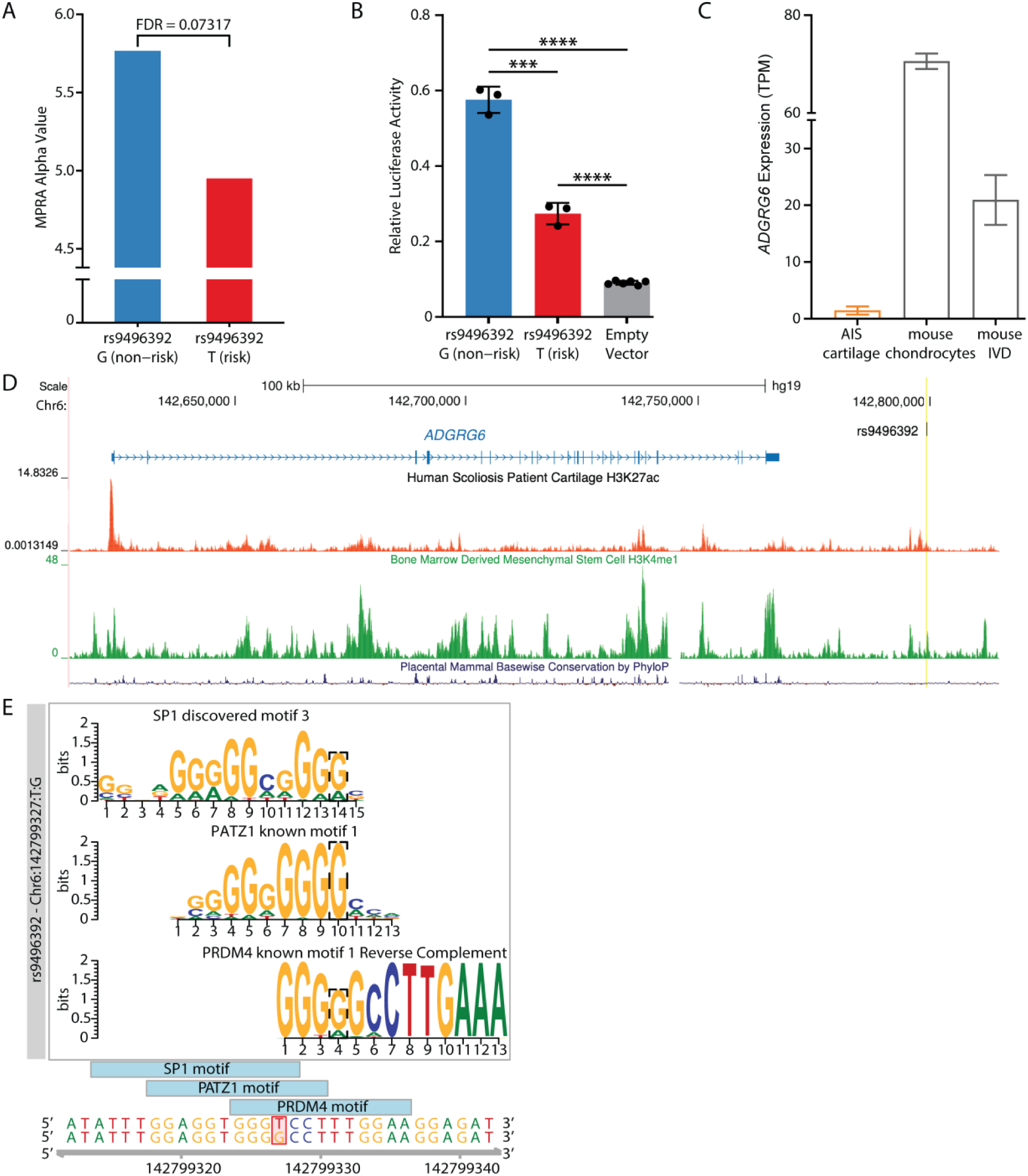
Predicted transcriptional disruption due to variant rs9496392. A) MPRA signal of rs9496392 CRS in TC28a2 (n=3). Reported FDR value is a false discovery rate adjusted likelihood ratio test *p*-value of the difference in activity between alleles reported by MPRAnalyze. B) Relative luciferase activity of the rs9496392 CRS in TC28a2 (n=3). ***, p < 0.001; ****, p < 0.0001. Statistical significance between CRS activity and empty-vector negative controls was determined by one-way ANOVA; statistical significance between SNP alleles was determined by unpaired Student’s *t*-test. C) Expression of *ADGRG6* in AIS spinal cartilage, mouse chondrocytes, and mouse intervertebral disc. D) Scoliosis H3K27ac ChIP-seq (red) and mesenchymal stem cell-derived chondrocyte H3K4me1 (green) enrichment at the rs9496392 enhancer (highlighted in yellow). E) Predicted disruption of transcription factor binding motifs. Error bars indicate mean ± standard deviation.

The rs9496392 variant is located approximately 32 kb downstream of the *ADGRG6* gene and coincides with H3K27ac enrichment in AIS patient cartilage (Makki et al. 2021) and H3K4me1 in mesenchymal stem cell-derived chondrocytes (Kundaje et al. 2015) (Roadmap Epigenomics Consortium ID: E049; Figure 4D). GTEx data identifies this SNP as an expression quantitative trait locus (eQTL) targeting the *ADGRG6* gene in lower leg skin, suprapubic skin, esophagus muscularis mucosa, gastrocnemius medialis, and subcutaneous adipose tissue (Dong et al. 2023; Boyle et al. 2012). As enhancers are tissue-dependent, and their target gene may vary between tissues, we wanted to confirm that rs9496392 interacts with *ADGRG6* in chondrocytes. Using published Hi-C data from the C28/I2 human chondrocyte cell line (Thulson et al. 2022), which has a similar gene expression profile to the TC28a2 cell line (Finger et al. 2003), we confirmed that the region flanking rs9496392 has some degree of interaction with the *ADGRG6* promoter (Supplemental Figure 10). We next wanted to determine potential upstream regulators of this enhancer. The genomic area encompassing rs9496392 displayed enrichment of several TFs by ChIP across several biosamples (Boyle et al. 2012), indicating that the enhancer may be bound by these TFs. ChIP-seq data showing peaks at rs9496392 was available for CTCF in 15 biosamples, MAFK in 5 biosamples, NFE2L2 in 3 biosamples, RAD21 in 2 biosamples, and several other transcription factors across various other biosamples (Supplemental Table 9). Interestingly, motifbreakR analysis predicts that the AIS risk allele of rs9496392 (T) strongly disrupts TFBSs for SP1, PATZ1, and PRDM4 (Figure 4E). These findings suggest that rs9496392 may affect expression of *ADGRG6* and other cartilage-related genes by altering transcription factor binding affinity at this enhancer.

## DISCUSSION

AIS is a common childhood disease that has significant impacts on quality of life, however little is known about the underlying etiology. GWAS have provided a glimpse into potential underlying mechanisms, though a drawback of these studies is their reliance on association, which leaves the contribution and function of associated variants as an open question. Few AIS regulatory variants have been identified thus far, as traditional individual reporter assays are low-throughput, time consuming, and are biased due to their reliance on correlative data to identify putative enhancers. Here we performed the first massively parallel assessment of AIS-associated variants to gain a better understanding of the genetic basis of AIS and the gene regulatory networks that may be perturbed in patients. To that end, we identified 1,664 variants in high linkage disequilibrium with 26 lead SNPs that were previously identified by GWAS and tested all possible alternate alleles to identify variants that differentially affect gene regulatory sequences in a high-throughput, unbiased manner.

In this work, we focused on identifying functional AIS variants that disrupt gene regulatory activity in chondrocytes, the primary cell type of cartilage, as cartilage has been widely implicated in AIS pathogenesis through studies in patients and model organisms (Ogura et al. 2015; Karner et al. 2015; Liu et al. 2021; Ushiki et al. 2024; C. A. Wise et al. 2020; H. Yu et al. 2024; Makki et al. 2021). We selected two chondrocyte cell lines that are commonly used in the field of cartilage biology, TC28a2 and SW1353. We identified 659 unique genomic positions active in TC28a2 and 499 unique genomic positions active in SW1353; of which, 886 were shared between both cell lines. This is the first functional identification of active enhancer elements in chondrocytes to be performed *en masse*, providing a new catalog of active enhancers for TC28a2 and SW1353.

Next, we identified CRSs by reference-alternate variant pairs to identify those with significant differences in transcription between the two alleles and with at least one allele being significantly active in the cell line. Our analysis identified a list of 234 highly confident variant pairs with allele-dependent regulatory activity, of which 92 are present in our selected HapMap populations. The remaining significant variants are not present in the selected populations but have similar numbers to the observed alternate alleles at each stage of the analysis. The 92 identified regulatory variants are spread across 18 of the 26 AIS GWAS loci. Some of these loci have a known role in AIS pathogenesis, such as *ADGRG6* and *PAX1*, and identification of novel functional AIS-associated variants at these loci will open many avenues for future investigations into the role these enhancer variants play in gene regulation. Other functional variants are at loci with unknown roles in AIS pathogenesis such as *CDH13* and *MIR4300HG*. Identification of functional variants at these loci will prompt new studies to investigate the mechanisms by which these loci affect AIS pathogenesis. Of the 92 regulatory variants, 79 are predicted to disrupt TFBS, which are critical for normal enhancer activity, and thus increases our understanding of potential mechanisms of effect for these variants. We attempted investigating the overlap of the 92 significant variants with markers for active enhancers, including ATAC-seq and H3K27ac ChIP-seq peaks, but as this data is not available for the two cell lines used in this study, we were unable to properly ascertain variant pairs overlapping ATAC or H3K27ac regions. Given the cell-specific context of epigenetic modifications, and that epigenetic data from chondrocytes is largely lacking, generation of these data sets will be an important step toward understanding the regulatory mechanisms governing disorders that affect cartilage. Future studies will be aimed at directly linking functional variants in enhancer elements to effects on transcription factor binding and to changes in target gene expression, thus elucidating gene regulatory networks underlying AIS pathogenesis.

Interestingly, one of the variants with differential regulatory activity in both cell lines (rs9496392) is located at the *ADGRG6* locus, which has a high relevance to cartilage and chondrocyte biology. In addition, the region flanking this SNP has chromatin modifications that are characteristic of enhancers (H3K27ac and H3K4me1) in cartilage and chondrocytes, is bound by several transcription factors in different tissues, and is predicted to have some interaction with the ADGRG6 promoter. *ADGRG6* encodes an adhesion G protein coupled receptor, previously called *GPR126*, that has established roles in chondrogenesis and chondrocyte homeostasis. Cartilage-specific knockout of *ADGRG6* in the osteochondral progenitor lineage and committed chondrocytes resulted in a postnatal scoliosis-like phenotype in mice that develops around the onset of puberty (Karner et al. 2015; Liu et al. 2021), while conditional deletion in osteoblasts resulted in reduced bone mineralization and body length (Sun et al. 2020). The novel regulatory variant (rs9496392) identified in our study is located downstream of the *ADGRG6* locus and is predicted to interact with the *ADGRG6* promoter. The risk allele of the variant is predicted to reduce affinity for several transcription factors including SP1, which has been shown to transactivate *COL2A1* expression (Chadjichristos et al. 2003; Ghayor et al. 2001). Recent studies have shown that ADGRG6 acts as an upstream regulator of COL2A1, among other ECM genes, in the postnatal mouse intervertebral disc. It is interesting to speculate that SP1 may regulate *ADGRG6*, and that its dysregulation in AIS may contribute to compromised ECM integrity (Aceves et al. 2026).

While this study identified a high-confidence set of AIS-associated regulatory variants with allele-dependent activity in chondrocyte-derived cells, a number of limitations warrant further investigation in future follow-up studies. First, due to the complexity and size of the variant library, at approximately 7 million associated barcodes with 735 average number of barcodes per CRS, we observed lower integration rates than anticipated. This lower representation may reduce sensitivity to detect regulatory effects, particularly for variants with modest allelic effects or lower barcode support, and may therefore increase the false-negative rate. However, the significant findings reported here are supported by multiple empirical reproducibility metrics, including strong CRS-level replicate concordance for aggregated DNA counts, RNA counts, and RNA/DNA ratios, as well as high agreement between full-barcode and barcode-subsampled MPRAnalyze results. Thus, while improved integration efficiency or expanded coverage may identify additional disruptive variants in future studies, these quality-control analyses support the robustness of the significant CRS-level findings reported here.

Second, although we have used genomic proximity to putatively annotate significantly disruptive variants with nearby genes, we recognize that proximity alone does not establish the affected gene or regulatory mechanism. Additional functional and mechanistic studies will be necessary to define the relevant downstream regulatory pathways.

To our knowledge, this is the first MPRA of AIS variants and the first MPRA to be performed in chondrocytes. This provides a ranked list of high-confidence functional AIS-associated variants with allele-specific regulatory ability, as well as a catalogue of active gene regulatory elements in chondrocytes. In this study, we focused on chondrocytes due to their known role in AIS pathogenesis through studies in patients and model organisms. Future studies will expand this analysis to other tissues relevant to AIS pathogenesis including bone, muscle, and sensory neurons. These findings will pave the way for the development of diagnostic and prognostic tests, as well as targeted therapies.

## METHODS

### MPRA library design

The MPRA design included 26 lead SNP positions identified in previous GWAS (C. A. Wise et al. 2020). We then expanded the library to include 1,638 additional variants that are in linkage disequilibrium (LD) greater than or equal to 0.7 with these lead SNPs in the CEU, CHB, and JPT HapMap3 populations (The 1000 Genomes Project Consortium et al. 2015). We included all four alleles for each of the SNPs, and only the reference and single alternate allele for indels. In total, the library includes 4,708 variant pairs (1,522 reference SNPs × 3 alternate alleles + 142 reference/alternate indels) creating a total of 6,372 tested CRSs (4,708 alternate alleles + 1,644 reference alleles). For positive controls, we selected 1,000 sequences centered on the 1,000 most significant q-valued H3K27ac ChIP-seq peaks identified in spinal cartilage (Makki et al. 2021) (SRA accession PRJNA625649) as identified by MACS2 (Zhang et al. 2008). Peaks were then filtered down to 601 that are within 1 mega-base of a transcription start site for a gene with at least 1 Transcript Per Million (TPM) RNA-seq value in either TC28a2 or SW1353 cell line. Negative controls were generated by randomly selecting and scrambling 200 of the reference allele candidate regulatory sequences. Each CRS was generated as a 200 base pair sequence with the allele of interest at base pair 101. For insertions and deletions (indels), base pair 101 is the first base of the added sequence.

### lentiMPRA library cloning and preparation of association sequencing library

The CRS library was cloned according to the protocol described by Gordon, Inoue, Martin, Schubach, et al (Gordon et al. 2020). In brief, the candidate regulatory sequence (CRS) library was synthesized by Agilent. Vector homology overhangs, a minimal promoter, sequencing adaptors, and 15-bp random barcodes were added to the CRSs by sequential rounds of PCR. CRS inserts were then cloned into linearized pLS-SceI (Addgene, cat no. 137725) using the NEBuilder HiFi DNA Assembly Master Mix (NEB, cat no. E2621S), electroporated into 10-beta electrocompetent cells (NEB, cat no. C3020K), and plated on carbenicillin selection plates overnight at 37°C. Colonies were pooled and plasmids were isolated by midiprep (Qiagen, cat no. 12945). To associate barcodes with CRSs, the CRS-barcode region was amplified by PCR using primers appended with Illumina flowcell adapters. Association sequencing was carried out on an Illumina NovaSeq X (PE150) with custom primers (Supplemental Table 10).

### Barcode-CRS association

DNA reads from the association library were processed using the MPRAflow pipeline (Gordon et al. 2020) and aligned to the CRSs using Bowtie 2 (Langmead and Salzberg 2012) with the “very sensitive” present parameters. Sequences were retained only if they contained an exact match to the CRS (--cigar 270M) and that barcode uniquely mapped to that CRS at least 70% of the time (--min-frac 0.70). Additionally, barcodes were associated with a CRS only if there were at least three instances of that barcode associating to that CRS (min-cov 3). We observed 30,673,678 barcodes across all association sequences and confidently associated 7,361,694 barcodes (24%) to CRSs at an average of 735 barcodes per CRS.

### Cell lines

The TC28a2 cell line was acquired from Dr. Miguel Otero, courtesy of Dr. Mary Goldring. SW1353 (cat no. HTB-94) and 293T (cat no. CRL-3216) cells were purchased from ATCC. All cells were cultured in DMEM (Life Technologies, cat no. 11995-065) supplemented with 10% fetal bovine serum (Corning, cat no. MT35010CV) and 1% penicillin-streptomycin (Life Technologies, cat no. 15140122) and maintained in a 5% CO_2_ humidified atmosphere.

### Lentivirus packaging

293T cells were co-transfected with the CRS library, pMD2.G (Addgene, cat no. 12259), and psPAX2 (Addgene, cat no. 12260) plasmids using the EndoFectin Lenti transfection reagent (GeneCopoeia, cat no. EF001) per the manufacturer’s protocol. The cell culture media was replaced with fresh media supplemented with 5% FBS, 1% penicillin-streptomycin, and 1× ViralBoost reagent (Alstem, cat no. VB100) after 8 hours. After 24 hours, media was collected, filtered through a 0.45 μm PES filter to remove cellular debris, and viral particles were concentrated using the Lenti-X concentrator reagent (Takara Bio, cat no. 631232). To determine virus titer, cells were seeded so as to yield 80% confluency the next day in a 24-well plate, then the media was replaced with DMEM supplemented with polybrene (Sigma-Aldrich, cat no. TR-1003-G), and increasing amounts of virus was added to each well (0, 1, 2, 4, 8, 16, 32, 64 μL). Media was changed after 24 hours and genomic DNA was extracted after 48 hours using the Wizard SV genomic DNA purification kit (Promega, cat no. A2361). Titer and MOI was determined by qPCR using the ΔCt method with primers targeting viral WPRE, plasmid backbone, and human LP34 genomic DNA.

### Lentiviral infection and preparation of barcode sequencing library

Lentivirus infection of TC28a2 and SW1353 cell lines were carried out in triplicate. Cells were seeded in 15-cm dishes so as to yield 80% confluency the next day (approximately 3.9 million TC28a2 cells or 2.7 million SW1353), then the following day media was replaced with media containing polybrene. To maintain library complexity, cells were transduced at a high Multiplicity of Infection (MOI ∼100), as determined by prior titration for optimal viability (Gordon et al. 2020). With a total of ∼3.9 × 10^6^ TC28a2) and ∼2.7 × 10^6^ (SW1353) cells per replicate and a functional titer of 0.5-1 × 10^6^ TU/µL, we achieved approximately 270–390 million integrations per sample. Based on the full barcode library complexity, this corresponds to an average representation of approximately 9–13 integrations per barcode across the input library. The next day fresh media was added, and 48 hours after infection DNA and RNA were simultaneously extracted using the AllPrep DNA/RNA mini kit (Qiagen, cat no. 80204). Cells were lysed in RLT Plus lysis buffer supplemented with 2-mercaptoethanol, scraped off the dish, and homogenized using a 3 mL syringe and 20-gauge needle. RNA was treated with the RNase-free DNase set (Qiagen, cat no. 79256) according to the manufacturer’s protocol, followed by treatment with the TURBO DNase kit (Life Technologies, cat no. AM1907) according to the manufacturer’s protocol for rigorous DNase treatment. RNA was then reverse transcribed using the SuperScript II reverse transcriptase (Life Technologies, cat no. 18064-071) and a primer downstream of the barcode containing an UMI and an Illumina P7 flow cell sequence.

To generate the barcode sequencing library, a P5 flowcell sequence and unique sample indexes were added to each DNA/RNA sample from each replicate by 3 cycles of PCR. Then, a second round of PCR was carried out with P5 and P7 primers. The number of PCR cycles for the second-round PCR was determined by qPCR; i.e. the cycle where the amplification curve nearly plateaus. The second-round PCR product was gel extracted and purified using the NucleoSpin Gel & PCR Clean-up mini kit (Macherey-Nagel, cat no. 740609.50) and quantified using a Qubit 3.0 fluorometer with the high sensitivity dsDNA kit (Life Technologies, cat no. Q33230). Finally, DNA and cDNA libraries were pooled in a 1:3 ratio before sequencing. Sequencing was performed on an Illumina NextSeq (PE75) with custom primers corresponding to the barcode, sample index, and UMI (Supplemental Table 10).

### Barcode counting

DNA and RNA reads from the assay were processed using the MPRAflow pipeline (Gordon et al. 2020) to obtain the number of barcode counts to be attributed to its associated CRSs. Reads were required to exactly match the barcode (bc-length 15). CRSs were retained for further analysis only if they contained at least 10 associated barcodes with observable DNA and RNA reads (--thresh 10; --merge_intersect TRUE). The TC28a2 cell line contained counts for 15,386,987 total barcodes, including 4,878,603 (66.27%) confidently assigned barcodes. The SW1353 cell line contained counts for 17,264,142 total barcodes, including 4,563,152 (61.98%) confidently assigned barcodes.

### Quantitative analysis of transcriptional activity

Quantification of a CRS transcriptional activity was calculated utilizing MPRAnalyze (Ashuach et al. 2019). Briefly, MPRAnalyze first calculates library wide normalization factors then fits generalized linear models (GLM) to the barcode counts. The program fits GLM to the latent construct estimates and observed DNA counts as well as fits a second nested GLM to the latent rate of transcription from the latent construct estimates and the observed RNA counts.

MPRAnalyze then optimizes the GLM’s via a gamma likelihood maximization of the DNA counts and a negative binomial likelihood maximization of the RNA counts. Transcriptional activity of the sequences is calculated utilizing the analyzeQuantification command to compare all sequences to the activity of the negative controls, while normalizing for library-wide and replicate variation. This outputs an estimated transcriptional “alpha” value, and a median absolute deviation (MAD) *Z*-score based *p*-value comparing the alpha value to the median alpha values of the negative control sequences. CRSs are reported as significantly active compared to negative controls with a *p*-value ≤ 0.05. Transcriptional activity variation between reference and alternate allele pairs are calculated utilizing the analyzeComparative command of MPRAnalyze.

This command provides a more sensitive calculation between the reference and alternate sequences than only comparing the alpha values from the analyzeQuantification command. This command includes the same normalizations as in the analyzeQuantification command but also incorporates the normalization of the different barcodes associated with the reference or alternate allele in the sequence pair. Additionally, the command normalized the overall difference between the reference and alternate allele to the RNA GLM to act as the null hypothesis. This command outputs the same alpha values as before but now the *p*-value is the significant difference between the reference and alternate allele, while accounting for each allele’s difference from the negative control. Alternate alleles have a significant difference compared to their reference allele sequence pair with a false discovery rate (FDR) ≤ 0.10. As the model fits the numerous barcode counts to the GLM models and reports the fitted alpha values, standard error metrics are not typically reported. Rather, they are incorporated into the *p*-values comparing the activity of the allele to either the negative controls or the other allele in the variant pair.

Due to the large complexity of the library, we were computationally unable to use all counted barcodes for the nested GLM calculation of differential allelic expression between variant pairs. To account for this, we selected 1 representative variant pair at each of the loci tested and generated the MPRAnalyze Comparative results on those variant pairs. We then subsetted the full variant pair table to only analyze 500 barcodes per CRS, which we were able to analyze with our available computational hardware. We then compared the results between the two approaches for variant pairs analyzed in both. We identified a 99.5% and 99.8% correlation of Log Fold Change between alleles for TC28a2 and SW1353, respectively, and a high correlation between the FDR values calculated, with only 1 out of a subset of variants pairs tested using all available barcodes, losing significance. With this high correlation of both Log Fold Change and FDR, we felt confident to continue the analysis, calculating differential allelic expression using only 500 barcodes per CRS.

To generate a final set of variants that have a high confidence of transcriptional disruption, we overlap the results of the analyzeQuantification and the analyzeComparison. For a variant pair to be considered high confidence, the alleles must be significantly different from each other as reported by the analyzeComparative analysis. To add confidence that the disruption is having an effect in the cell line, we also require the variant pair to have at least one of the two alleles be significantly active compared to the negative controls in the analyzeQuantification analysis. This prevents any variant pairs that are significantly different from each other, but neither have an effect in the cell line as the transcription rates are at the same level as the negative controls.

### Luciferase validation assays

CRSs (Supplemental Table 11) were amplified from human genomic DNA (Promega, cat no. G304A) by PCR and cloned into pGL4.23[luc2/mP] (Promega, cat no. E8411) using the NEBuilder HiFi DNA assembly kit (NEB, cat no. E2621S), then transformed into DH5-alpha competent cells (NEB, cat no. C2987H)) and plated on carbenicillin selection plates overnight at 37°C. Successful insertion of the CRS was verified by colony PCR, plasmids were isolated by miniprep (Macherey-Nagel, cat no. 740490.50), and sequenced. The alternate SNP alleles were introduced by site-directed mutagenesis using the Phusion Site-Directed Mutagenesis kit (ThermoFisher, cat no. F541). Cell lines were plated in 24-well plates to yield 80% confluency the next day, then transfected with 900 ng of the CRS plasmid and 100 ng of pGL4.74[hR/luc] (Promega, cat no. E6921) using X-tremeGENE HP DNA Transfection Reagent (Roche, cat no. 6366244001) according to the manufacturer’s protocol. After 48 hours post-transfection, luciferase activity was assayed using the Dual Luciferase Reporter Assay System (Promega, cat no. E1980) according to the manufacturer’s protocol on a Glomax Multi Detection System plate reader (Promega, cat no. 9301-010).

CRS activity was measured in comparison to cells transfected with empty vector negative control plasmids (pGL4.23) with at least 6 biological replicates for each construct. Statistical significance between CRSs and negative controls was calculated by one-way ANOVA with Dunnett’s test to account for multiple comparisons, while statistical significance between SNP alleles of the same CRS was calculated by unpaired *t*-test with Benjamini-Krieger-Yekutieli correction for multiple comparisons.

### Transcription factor binding disruption predictions

Transcription factor binding site disruptions were predicted using the program MotifBreakR (Coetzee et al. 2015). Briefly, MotifBreakR utilizes a position weight matrix from the ENCODE database of transcription factor binding sites to predict if a variant causes a neutral, weak, or strong disruption to a predicted binding site. In our study we only included disruptions to binding sites if the significance *p*-value of disruption was smaller than or equal to 1×10-4 and the disruption to the binding site was listed as weak or strong. For our analysis we further filtered any binding site that was for a transcription factor not expressed (< 1 TPM) in the cell line that the variant pair was found significant in.

### Generation of TC28a2 RNA-seq data and processing of TC28a2 and SW1353 RNA-seq

Total RNA from the TC28a2 cell line was isolated using the Qiagen RNeasy mini kit and mRNA was selected for using poly(T) oligo-attached magnetic beads. First strand cDNA was synthesized using random hexamers, followed by second strand cDNA synthesis. Paired-end sequencing was carried out on an Illumina NovaSeq6000. Raw RNA-seq FASTQ files for the SW1353 cell line were downloaded from the GEO database under accession number GSE176234.

For processing the sequenced FASTQ files, we utilized nf-core’s standardized RNA-seq pipeline (Version 3.12.0) (Ewels et al. 2020) with the following additional parameters. For the Trim Galore! step we include the following on top of the default parameters: –trim-n to remove trailing N’s on either side of the read; –length 40 to remove reads with a trimmed size less than 40 bases; –quality 20 to remove ends of reads with below 20 Phred scores; –clip_R1 5 and clip_R2 5 to trim the first 5 bases off of the 5’ and 3’ end of the reads. Additionally, we include – min_trimmed_reads 5000 to remove any samples with below 5000 reads after trimming. We then include –skip_alignment, –skip_deseq2_qc, and –psuedo_aligner salmon with an igenomes reference file set to call transcript per million (TPM) counts for genes in the cell lines. The generated file is a two-column table with HGNC gene symbols and the TPM value. We then remove any gene symbols with a TPM value less than 1 to get a list of genes active in the SW1353 and TC28a2 cell lines respectively.

## DATA ACCESS

All raw and processed sequencing data generated in this study have been submitted to the NCBI Gene Expression Omnibus (GEO; https://www.ncbi.nlm.nih.gov/geo/) under accession number GSE316848 (MPRA) and GSE316849 (TC28a2 RNA-seq).

## ACKNOWLEDGMENTS

This work was funded by the Scoliosis Research Society (SRS) and the University of Florida Research Opportunity Seed Fund awarded to N.M.

## Author contributions

N.M., A.K., D.R., and J.K. conceived the experiments. D.R., F.R.T., and C.C. performed experiments. J.K., D.K., N.M., and A.K. analyzed the data. D.R., J.K., N.M., and A.K. wrote the manuscript.

