## Supplemental Figures and Tables for "Massively parallel characterization of adolescent idiopathic scoliosis risk variants": Supplemental_FiguresAndTables_updated references_no table legends.pdf

**Supplemental Tables**

**Supplemental Table 1.** List of lead SNPs and number of SNPs in LD.

| Lead SNP | Nearest Gene | Reference | Annotation | # Linked SNPs Tested |
|---|---|---|---|---|
| rs11190870 | *LBX1* | Haller et al. 2016 | intergenic | 13 |
| rs6570507 | *ADGRG6* | Kou et al. 2013 | intron | 81 |
| rs3904778 | *BNC2* | Ogura et al. 2015 | intron | 60 |
| rs6137473 | *PAX1* | Sharma et al. 2015 | intergenic | 269 |
| rs687621 | *ABO* | Khanshour et al. 2018 | intron | 47 |
| rs1455114 | *SOX6* | Khanshour et al. 2018 | intron | 74 |
| rs4513093 | *CDH13* | Khanshour et al. 2018 | intron | 138 |
| rs4940576 | *BCL2* | Zhu et al. 2015 | intron | 35 |
| rs13398147 | *PAX3/EPHA4* | Zhu et al. 2015 | intergenic | 50 |
| rs421215 | *AJAP1* | Zhu et al. 2015 | intergenic | 3 |
| rs12946942 | *SOX9, KCNJ2* | Miyake et al. 2013 | intergenic | 3 |
| rs35333564 | *MIR4300HG* | Ogura et al. 2017 | intron | 229 |
| rs141903557 | *LINC02994* | Kou et al. 2019 | intron | 124 |
| rs11205303 | *MTMR11* | Kou et al. 2019 | exon | 4 |
| rs12029076 | *ARF1* | Kou et al. 2019 | intron | 14 |
| rs1978060 | *TBX1* | Kou et al. 2019 | intron | 8 |
| rs2467146 | *LINC02378/MIR3974* | Kou et al. 2019 | intergenic | 99 |
| rs11787412 | *CSMD1* | Kou et al. 2019 | intron | 103 |
| rs188915802 | *KIF24* | Kou et al. 2019 | intron | 10 |
| rs658839 | *BCKDHB* | Kou et al. 2019 | intergenic | 60 |
| rs160335 | *CREB5* | Kou et al. 2019 | intron | 25 |
| rs482012 | *NT5DC1* | Kou et al. 2019 | intron | 32 |
| rs11341092 | *UNCX* | Kou et al. 2019 | intergenic | 49 |
| rs17011903 | *PLXNA2* | Kou et al. 2019 | intron | 37 |
| rs397948882 | *MEOX2* | Kou et al. 2019 | intergenic | 0 |
| rs12149832 | *FTO* | Kou et al. 2019 | intron | 97 |

**Supplemental Table 8.** List of genes that have been linked with AIS.

| Gene | Reference |
|---|---|
| *UNCX* | Yonezawa et al. 2020 |
| *EGR1* | Yonezawa et al. 2020 |
| *ADGRG6* | Karner et al. 2015; Liu et al. 2021 |
| *COL11A1* | Yu et al. 2024 |
| *PAX1* | Sharma et al. 2015; Yu et al. 2024 |
| *MMP3* | Yu et al. 2024 |
| *COL11A2* | Rebello et al. 2023 |
| *COL2A1* | Barat-Houari et al. 2016; Hoornaert et al. 2010 |
| *COL8A1A* | Gray et al. 2014 |
| *SLC39A8* | Haller et al. 2018 |
| *PTK7* | Hayes et al. 2014; Su et al. 2021 |
| *LBX1* | Wang et al. 2022; Decourtye et al. 2022 |
| *SOX6* | Smits et al. 2003 |
| *SOX9* | Henry et al. 2012 |
| *FBN1* | Buchan et al. 2014; de Azevedo et al. 2022 |
| *FBN2* | Buchan et al. 2014 |
| *FTO* | Wang et al. 2023 |
| *BNC2* | Ogura et al. 2015 |
| *MIR4300HG* | Ogura et al. 2017 |
| *C21ORF59* | Jaffe et al. 2016 |
| *CCDC40* | Becker-Heck et al. 2011; Xu et al. 2025 |
| *CCDC151* | Grimes et al. 2016 |
| *DYX1C1* | Bieder et al. 2023 |
| *KIF6* | Buchan et al. 2014 |

### Supplemental Figures

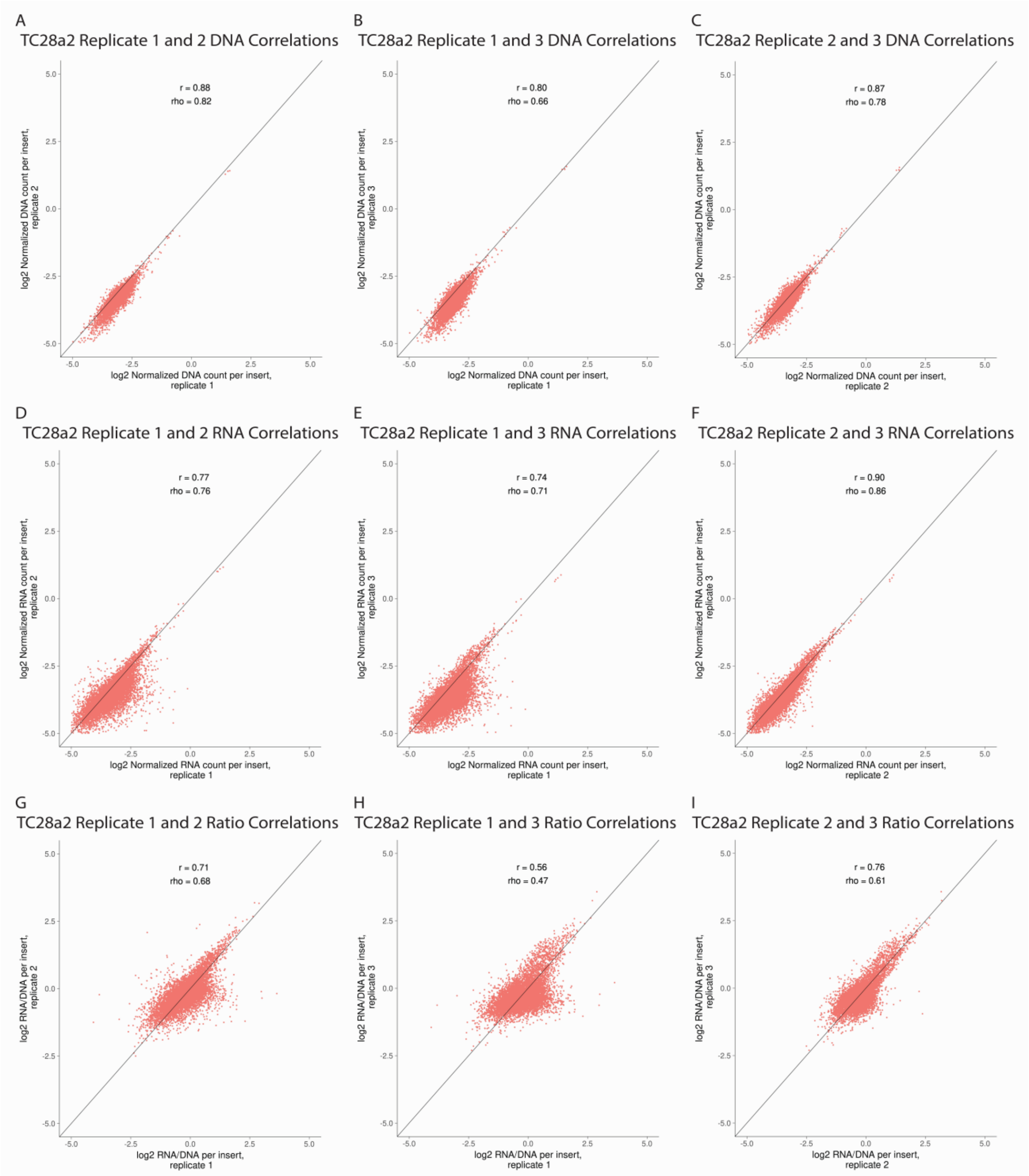


**Supplemental Figure 1.** TC28a2 DNA and RNA correlation across replicates. This figure shows the correlation of DNA (A-C), RNA (D-F), and RNA/DNA ratio (G-I) between the 3 replicates. Panels A, D, and G compare replicate 1 on the x-axis with replicate 2 on the y-axis. Panels B, E, and H compare replicate 1 on the x-axis with replicate 3 on the y-axis. Panels C, F, and I compare replicate 2 on the x-axis with replicate 3 on the y-axis.

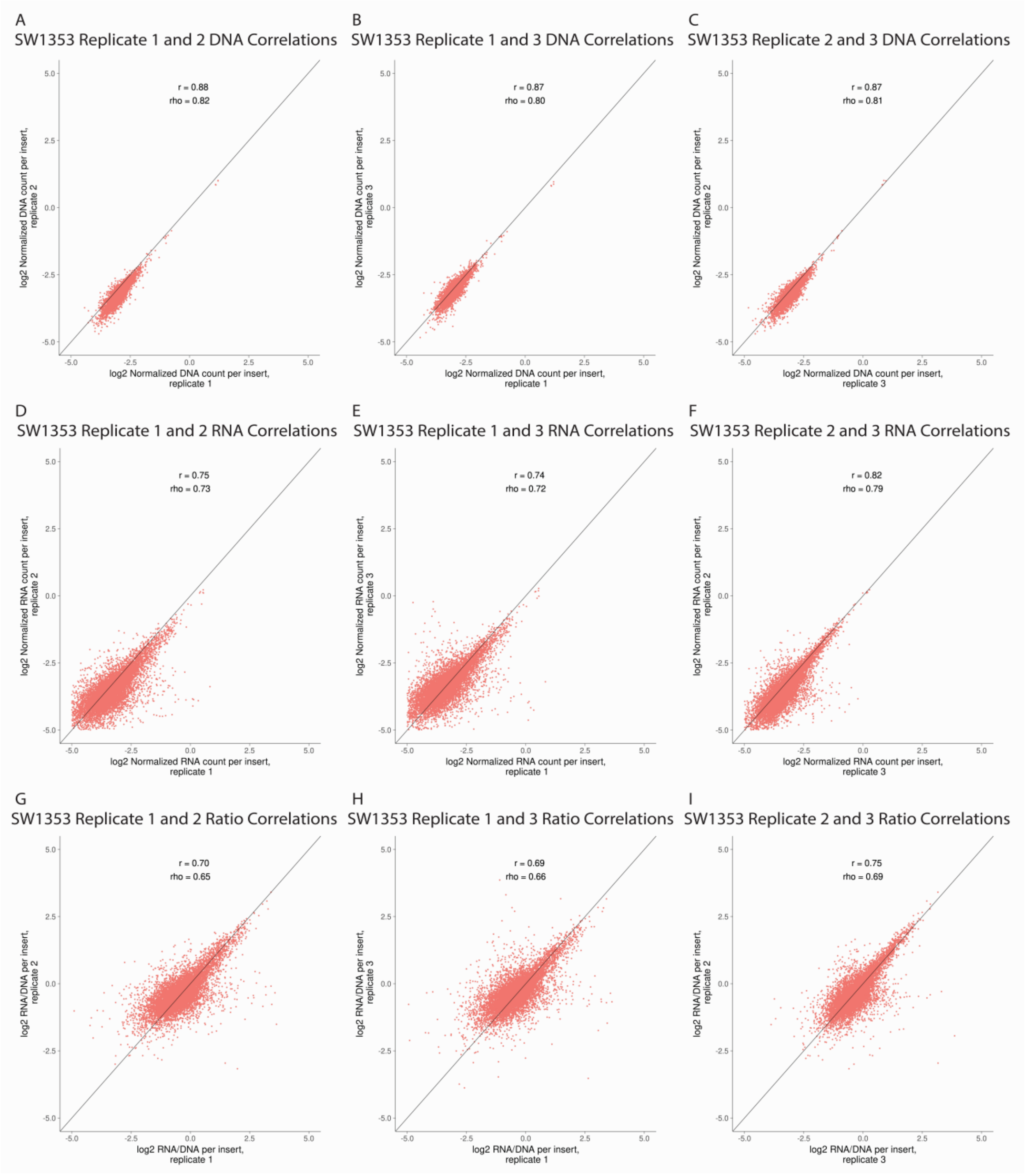


**Supplemental Figure 2.** SW1353 DNA and RNA correlation across replicates. This figure shows the correlation of DNA (A-C), RNA (D-F), and RNA/DNA ratio (G-I) between the 3 replicates. Panels A, D, and G compare replicate 1 on the x-axis with replicate 2 on the y-axis. Panels B, E, and H compare replicate 1 on the x-axis with replicate 3 on the y-axis. Panels C, F, and I compare replicate 2 on the x-axis with replicate 3 on the y-axis.

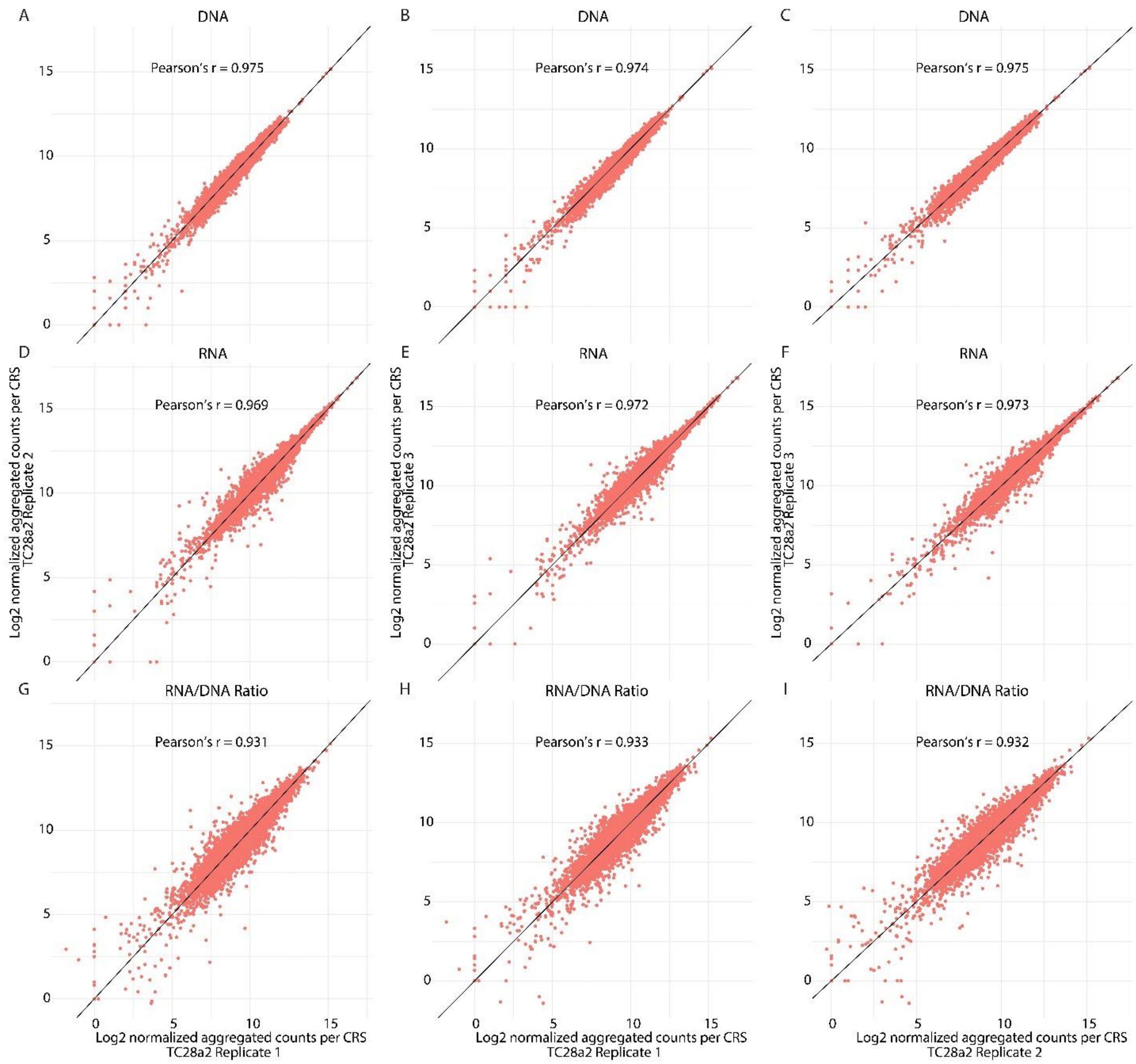


**Supplemental Figure 3.** TC28a2 CRS aggregated DNA and RNA correlation across replicates. This figure shows the correlation of DNA (A-C), RNA (D-F), and RNA/DNA ratio (G-I) between the 3 replicates after aggregating the counts for all barcodes associated with each CRS. Panels A, D, and G compare replicate 1 on the x-axis with replicate 2 on the y-axis. Panels B, E, and H compare replicate 1 on the x-axis with replicate 3 on the y-axis. Panels C, F, and I compare replicate 2 on the x-axis with replicate 3 on the y-axis.

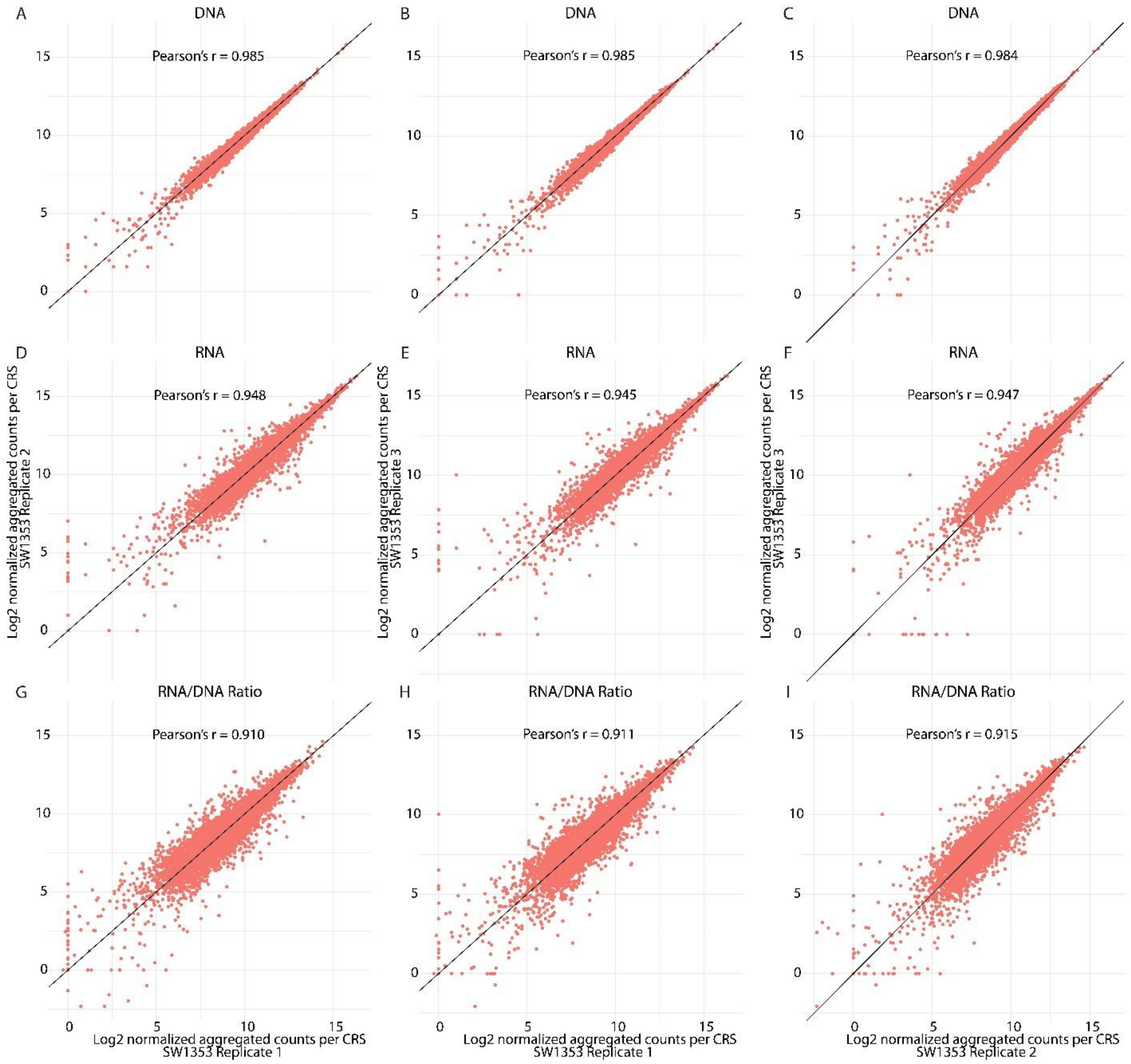


**Supplemental Figure 4.** SW1353 CRS aggregated DNA and RNA correlation across replicates. This figure shows the correlation of DNA (A-C), RNA (D-F), and RNA/DNA ratio (G-I) between the 3 replicates after aggregating the counts for all barcodes associated with each CRS. Panels A, D, and G compare replicate 1 on the x-axis with replicate 2 on the y-axis. Panels B, E, and H compare replicate 1 on the x-axis with replicate 3 on the y-axis. Panels C, F, and I compare replicate 2 on the x-axis with replicate 3 on the y-axis.

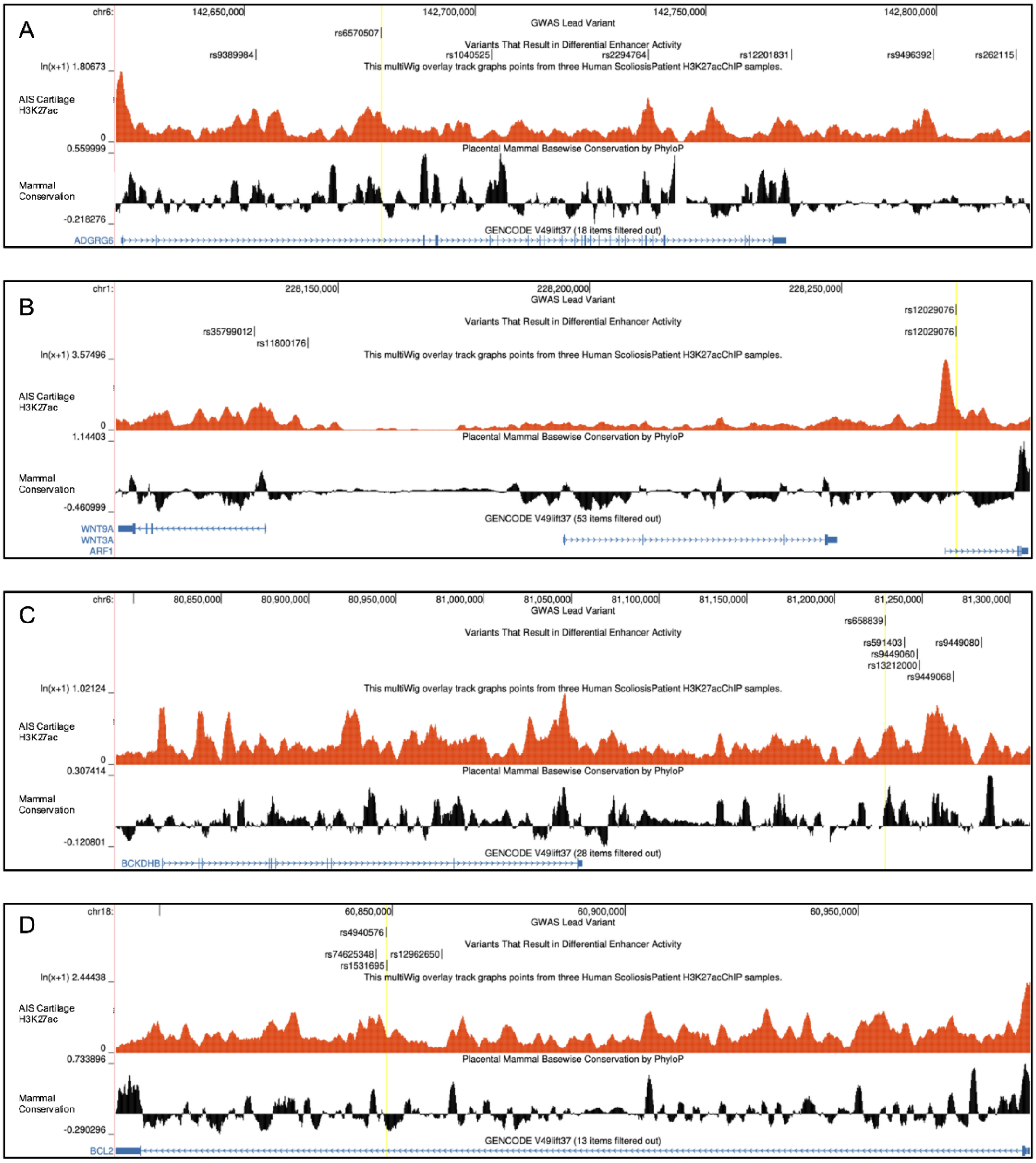

A
chr6:
142,650,000
142,700,000
142,750,000
142,800,000
GWAS Lead Variant
rs6570507
Variants That Result in Differential Enhancer Activity
rs9389984
rs1040525
rs2294764
rs12201831
rs9496392
rs262115
ln(x+1) 1.80673
This multiWig overlay track graphs points from three Human ScoliosisPatient H3K27acChIP samples.
AIS Cartilage H3K27ac
0
0.559999
Placental Mammal Basewise Conservation by PhyloP
Mammal Conservation
-0.218276
GENCODE V49lift37 (18 items filtered out)
ADGRG6
B
chr1:
228,150,000
228,200,000
228,250,000
GWAS Lead Variant
rs12029076
Variants That Result in Differential Enhancer Activity
rs35799012
rs11800176
rs12029076
ln(x+1) 3.57496
This multiWig overlay track graphs points from three Human ScoliosisPatient H3K27acChIP samples.
AIS Cartilage H3K27ac
0
1.14403
Placental Mammal Basewise Conservation by PhyloP
Mammal Conservation
-0.460999
GENCODE V49lift37 (53 items filtered out)
WNT9A
WNT3A
ARF1
C
chr6:
80,850,000
80,900,000
80,950,000
81,000,000
81,050,000
81,100,000
81,150,000
81,200,000
81,250,000
81,300,000
GWAS Lead Variant
rs658839
Variants That Result in Differential Enhancer Activity
rs591403
rs9449080
rs9449060
rs13212000
rs9449068
ln(x+1) 1.02124
This multiWig overlay track graphs points from three Human ScoliosisPatient H3K27acChIP samples.
AIS Cartilage H3K27ac
0
0.307414
Placental Mammal Basewise Conservation by PhyloP
Mammal Conservation
-0.120801
GENCODE V49lift37 (28 items filtered out)
BCKDHB
D
chr18:
60,850,000
60,900,000
60,950,000
GWAS Lead Variant
rs4940576
Variants That Result in Differential Enhancer Activity
rs74625348
rs12962650
rs1531695
ln(x+1) 2.44438
This multiWig overlay track graphs points from three Human ScoliosisPatient H3K27acChIP samples.
AIS Cartilage H3K27ac
0
0.733896
Placental Mammal Basewise Conservation by PhyloP
Mammal Conservation
-0.290296
GENCODE V49lift37 (13 items filtered out)
BCL2

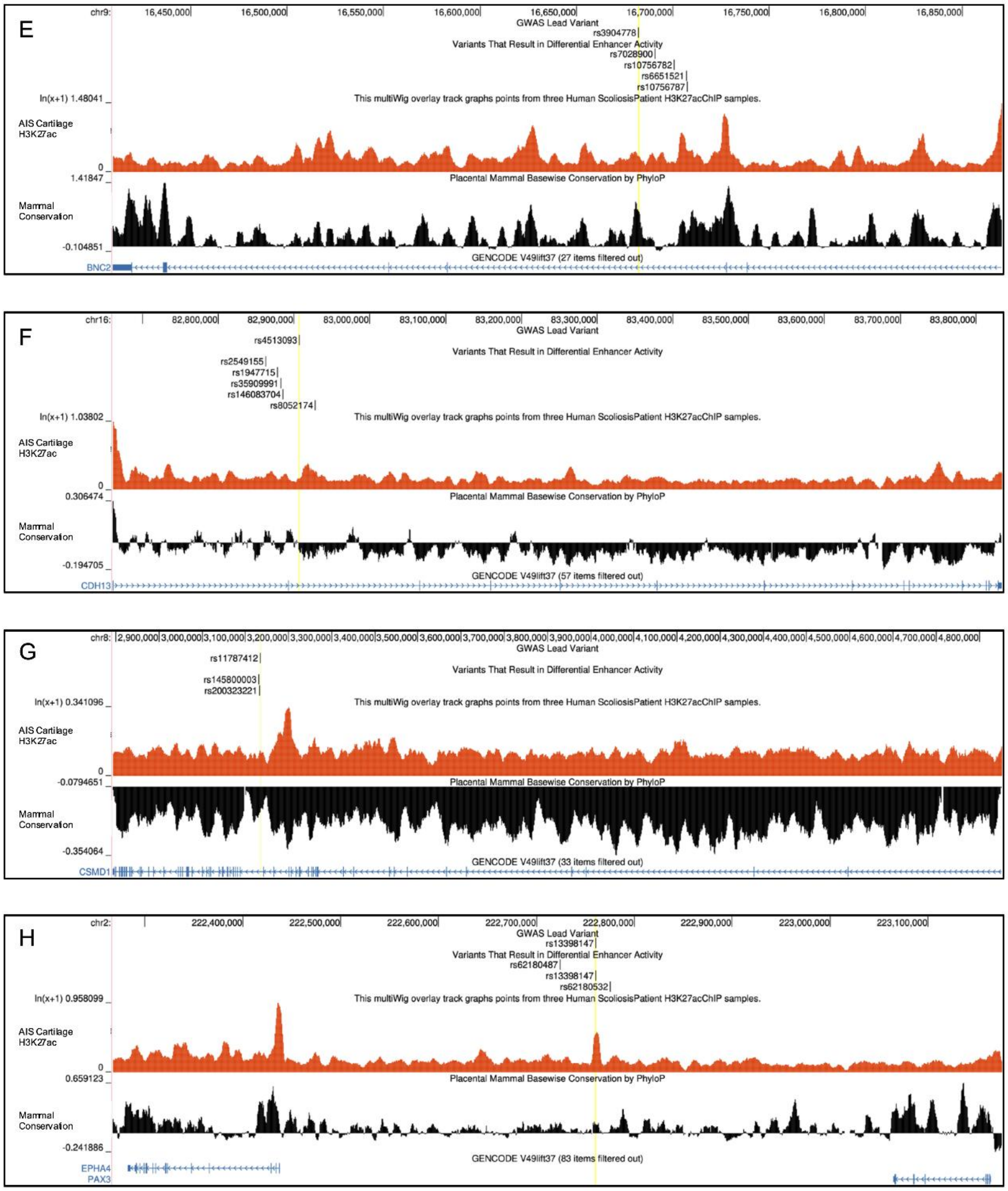
E
chr9:
16,450,000
16,500,000
16,550,000
16,600,000
16,650,000
16,700,000
16,750,000
16,800,000
16,850,000
GWAS Lead Variant
rs3904778
Variants That Result in Differential Enhancer Activity
rs7028900
rs10756782
rs6651521
rs10756787
ln(x+1) 1.48041
This multiWig overlay track graphs points from three Human ScoliosisPatient H3K27acChIP samples.
AIS Cartilage H3K27ac
0
1.41847
Placental Mammal Basewise Conservation by PhyloP
Mammal Conservation
-0.104851
GENCODE V49lift37 (27 items filtered out)
BNC2
F
chr16:
82,800,000
82,900,000
83,000,000
83,100,000
83,200,000
83,300,000
83,400,000
83,500,000
83,600,000
83,700,000
83,800,000
GWAS Lead Variant
rs4513093
Variants That Result in Differential Enhancer Activity
rs2549155
rs1947715
rs35909991
rs146083704
rs8052174
ln(x+1) 1.03802
This multiWig overlay track graphs points from three Human ScoliosisPatient H3K27acChIP samples.
AIS Cartilage H3K27ac
0
0.306474
Placental Mammal Basewise Conservation by PhyloP
Mammal Conservation
-0.194705
GENCODE V49lift37 (57 items filtered out)
CDH13
G
chr8:
2,900,000
3,000,000
3,100,000
3,200,000
3,300,000
3,400,000
3,500,000
3,600,000
3,700,000
3,800,000
3,900,000
4,000,000
4,100,000
4,200,000
4,300,000
4,400,000
4,500,000
4,600,000
4,700,000
4,800,000
GWAS Lead Variant
rs11787412
Variants That Result in Differential Enhancer Activity
rs145800003
rs200323221
ln(x+1) 0.341096
This multiWig overlay track graphs points from three Human ScoliosisPatient H3K27acChIP samples.
AIS Cartilage H3K27ac
0
-0.0794651
Placental Mammal Basewise Conservation by PhyloP
Mammal Conservation
-0.354064
GENCODE V49lift37 (33 items filtered out)
CSMD1
H
chr2:
222,400,000
222,500,000
222,600,000
222,700,000
222,800,000
222,900,000
223,000,000
223,100,000
GWAS Lead Variant
rs13398147
Variants That Result in Differential Enhancer Activity
rs62180487
rs13398147
rs62180532
ln(x+1) 0.958099
This multiWig overlay track graphs points from three Human ScoliosisPatient H3K27acChIP samples.
AIS Cartilage H3K27ac
0
0.659123
Placental Mammal Basewise Conservation by PhyloP
Mammal Conservation
-0.241886
GENCODE V49lift37 (83 items filtered out)
EPHA4
PAX3

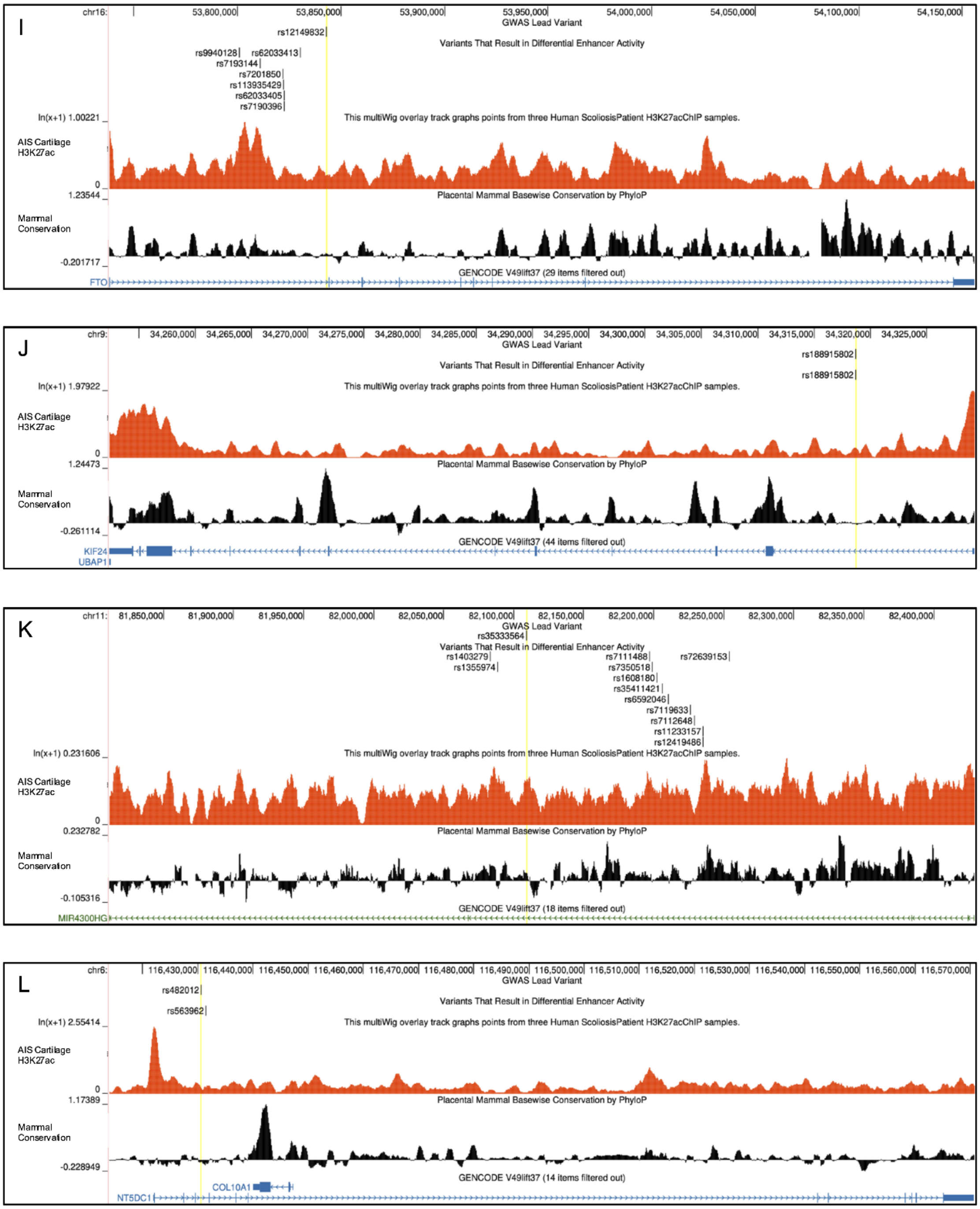

I
chr16:
53,800,000
53,850,000
53,900,000
53,950,000
54,000,000
54,050,000
54,100,000
54,150,000
GWAS Lead Variant
rs12149832
Variants That Result in Differential Enhancer Activity
rs9940128
rs62033413
rs7193144
rs7201850
rs113935429
rs62033405
rs7190396
ln(x+1) 1.00221
This multiWig overlay track graphs points from three Human ScoliosisPatient H3K27acChIP samples.
AIS Cartilage H3K27ac
0
1.23544
Placental Mammal Basewise Conservation by PhyloP
Mammal Conservation
-0.201717
GENCODE V49lift37 (29 items filtered out)
FTO
J
chr9:
34,260,000
34,265,000
34,270,000
34,275,000
34,280,000
34,285,000
34,290,000
34,295,000
34,300,000
34,305,000
34,310,000
34,315,000
34,320,000
34,325,000
GWAS Lead Variant
rs188915802
Variants That Result in Differential Enhancer Activity
rs188915802
ln(x+1) 1.97922
This multiWig overlay track graphs points from three Human ScoliosisPatient H3K27acChIP samples.
AIS Cartilage H3K27ac
0
1.24473
Placental Mammal Basewise Conservation by PhyloP
Mammal Conservation
-0.261114
GENCODE V49lift37 (44 items filtered out)
KIF24
UBAP1
K
chr11:
81,850,000
81,900,000
81,950,000
82,000,000
82,050,000
82,100,000
82,150,000
82,200,000
82,250,000
82,300,000
82,350,000
82,400,000
GWAS Lead Variant
rs35333564
Variants That Result in Differential Enhancer Activity
rs1403279
rs1355974
rs7111488
rs72639153
rs7350518
rs1608180
rs35411421
rs6592046
rs7119633
rs7112648
rs11233157
rs12419486
ln(x+1) 0.231606
This multiWig overlay track graphs points from three Human ScoliosisPatient H3K27acChIP samples.
AIS Cartilage H3K27ac
0
0.232782
Placental Mammal Basewise Conservation by PhyloP
Mammal Conservation
-0.105316
GENCODE V49lift37 (18 items filtered out)
MIR4300HG
L
chr6:
116,430,000
116,440,000
116,450,000
116,460,000
116,470,000
116,480,000
116,490,000
116,500,000
116,510,000
116,520,000
116,530,000
116,540,000
116,550,000
116,560,000
116,570,000
GWAS Lead Variant
rs482012
Variants That Result in Differential Enhancer Activity
rs563962
ln(x+1) 2.55414
This multiWig overlay track graphs points from three Human ScoliosisPatient H3K27acChIP samples.
AIS Cartilage H3K27ac
0
1.17389
Placental Mammal Basewise Conservation by PhyloP
Mammal Conservation
-0.228949
GENCODE V49lift37 (14 items filtered out)
COL10A1
NT5DC1

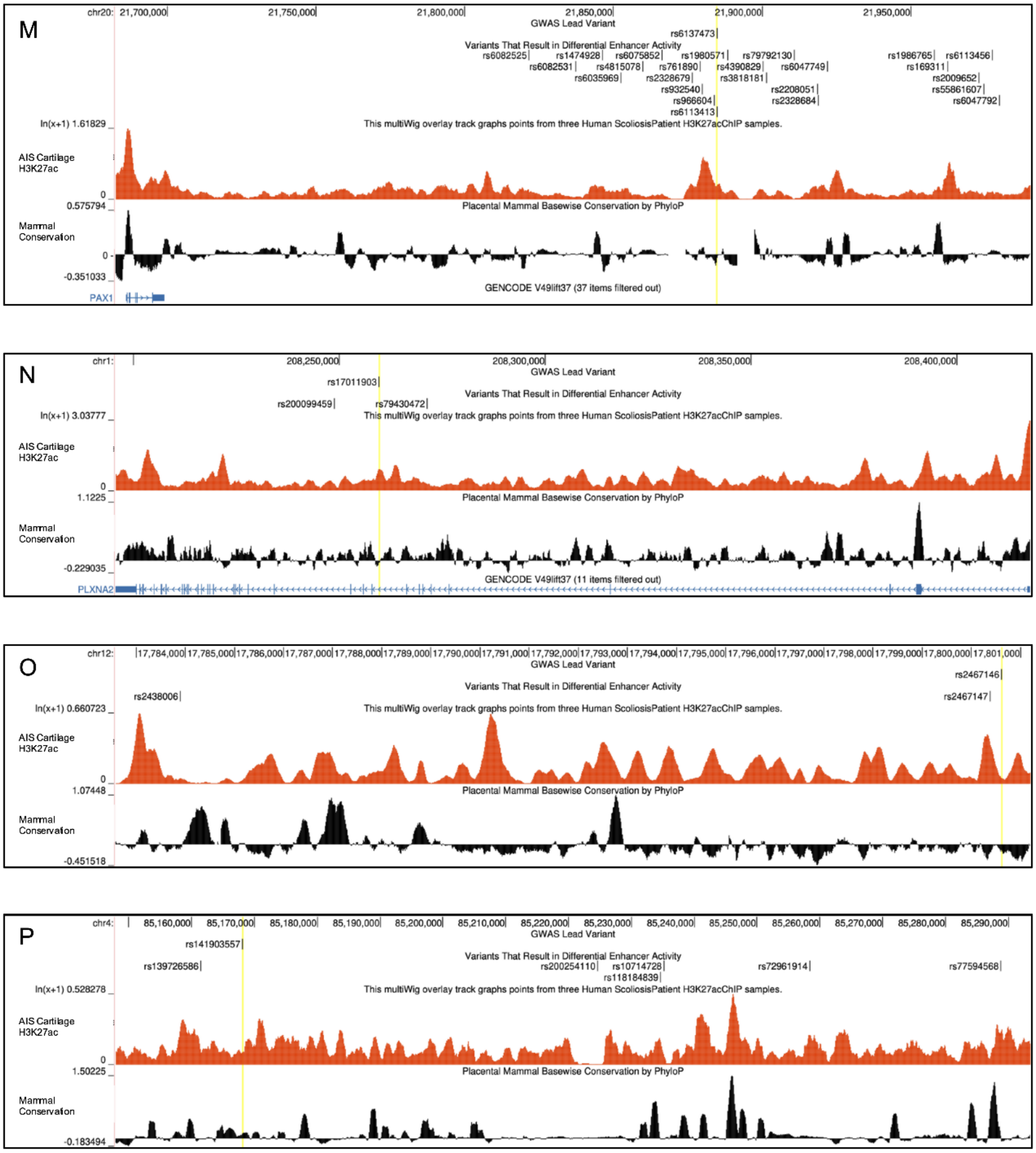

M
chr20: 21,700,000 21,750,000 21,800,000 21,850,000 21,900,000 21,950,000
GWAS Lead Variant
rs6137473
Variants That Result in Differential Enhancer Activity
rs6082525 rs1474928 rs6075852 rs1980571 rs79792130 rs1986765 rs6113456
rs6082531 rs4815078 rs761890 rs4390829 rs6047749 rs169311
rs6035969 rs2328679 rs3818181 rs2009652
rs932540 rs2208051 rs55861607
rs966604 rs2328684 rs6047792
rs6113413
ln(x+1) 1.61829
This multiWig overlay track graphs points from three Human ScoliosisPatient H3K27acChIP samples.
AIS Cartilage H3K27ac
0
0.575794
Placental Mammal Basewise Conservation by PhyloP
Mammal Conservation
0
-0.351033
GENCODE V49lift37 (37 items filtered out)
PAX1
N
chr1: 208,250,000 208,300,000 208,350,000 208,400,000
GWAS Lead Variant
rs17011903
Variants That Result in Differential Enhancer Activity
rs200099459 rs79430472
ln(x+1) 3.03777
This multiWig overlay track graphs points from three Human ScoliosisPatient H3K27acChIP samples.
AIS Cartilage H3K27ac
0
1.1225
Placental Mammal Basewise Conservation by PhyloP
Mammal Conservation
-0.229035
GENCODE V49lift37 (11 items filtered out)
PLXNA2
O
chr12: 17,784,000 17,785,000 17,786,000 17,787,000 17,788,000 17,789,000 17,790,000 17,791,000 17,792,000 17,793,000 17,794,000 17,795,000 17,796,000 17,797,000 17,798,000 17,799,000 17,800,000 17,801,000
GWAS Lead Variant
rs2467146
Variants That Result in Differential Enhancer Activity
rs2438006 rs2467147
ln(x+1) 0.660723
This multiWig overlay track graphs points from three Human ScoliosisPatient H3K27acChIP samples.
AIS Cartilage H3K27ac
0
1.07448
Placental Mammal Basewise Conservation by PhyloP
Mammal Conservation
-0.451518
P
chr4: 85,160,000 85,170,000 85,180,000 85,190,000 85,200,000 85,210,000 85,220,000 85,230,000 85,240,000 85,250,000 85,260,000 85,270,000 85,280,000 85,290,000
GWAS Lead Variant
rs141903557
Variants That Result in Differential Enhancer Activity
rs139726586 rs200254110 rs10714728 rs72961914 rs77594568
rs118184839
ln(x+1) 0.528278
This multiWig overlay track graphs points from three Human ScoliosisPatient H3K27acChIP samples.
AIS Cartilage H3K27ac
0
1.50225
Placental Mammal Basewise Conservation by PhyloP
Mammal Conservation
-0.183494

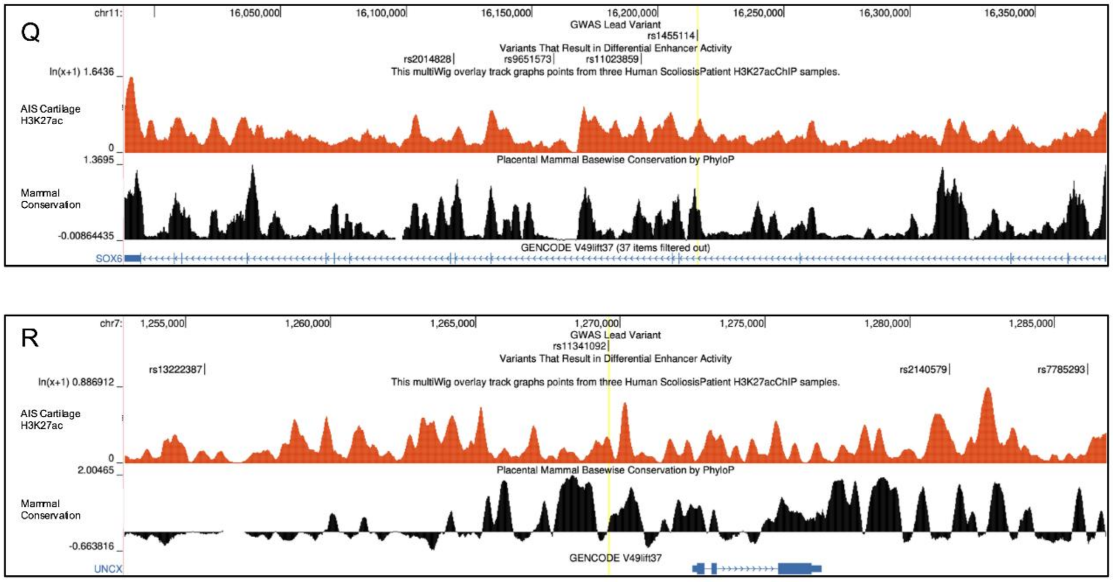


**Supplemental Figure 5.** UCSC Genome Browser view of significant regulatory variants in LD with GWAS lead variants (highlighted in yellow) overlapping with H3K27ac enrichment in AIS spinal cartilage (red, from Makki et al. (2020)), and base-wise conservation in placental mammals (black). A) Significant regulatory variants in LD with rs6570507. B) Significant regulatory variants in LD with rs12029076. C) Significant regulatory variants in LD with rs658839. D) Significant regulatory variants in LD with rs4940576. E) Significant regulatory variants in LD with rs3904778. F) Significant regulatory variants in LD with rs4513093. G) Significant regulatory variants in LD with rs11787412. H) Significant regulatory variants in LD with rs13398147. I) Significant regulatory variants in LD with rs12149832. J) Significant regulatory variants in LD with rs188915802. K) Significant regulatory variants in LD with rs35333564. L) Significant regulatory variants in LD with rs482012. M) Significant regulatory variants in LD with rs6137473. N) Significant regulatory variants in LD with rs17011903. O) Significant regulatory variants in LD with rs2467146. P) Significant regulatory variants in LD with rs141903557. Q) Significant regulatory variants in LD with rs1455114. R) Significant regulatory variants in LD with rs11341092.

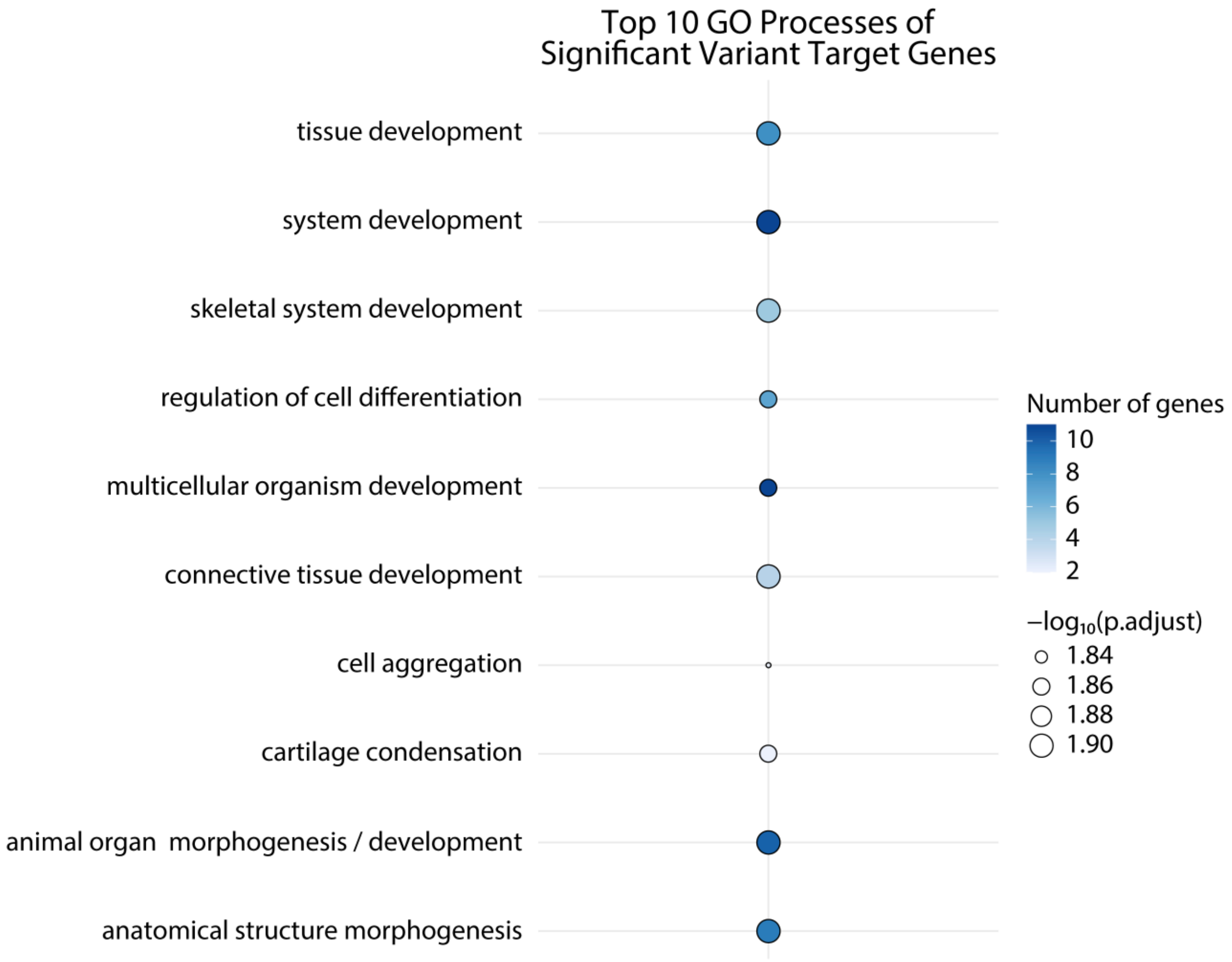


**Supplemental Figure 6.** Enriched biological processes of single nearest gene to each significant variant pair identified in either cell line using Gene Ontology (GO) analysis.

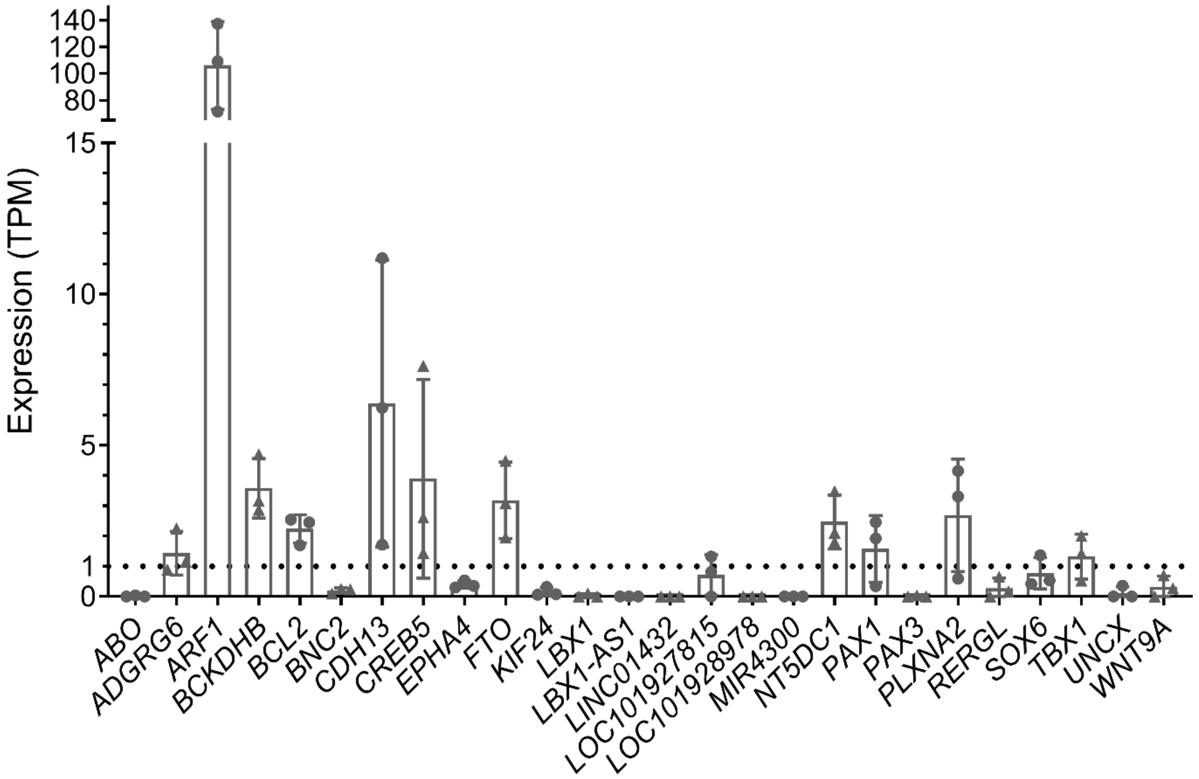


**Supplemental Figure 7.** Expression in TPM of genes proximal to variants with significant differential MPRA activity in human spinal cartilage. Notably ARF1, EPHA4, LOC153910, and WNT9A are only found proximal to significant main allele variants. While ABO, CREB5, LBX1-AS1, and TBX1 are only found proximal to significant synthetic allele variants.

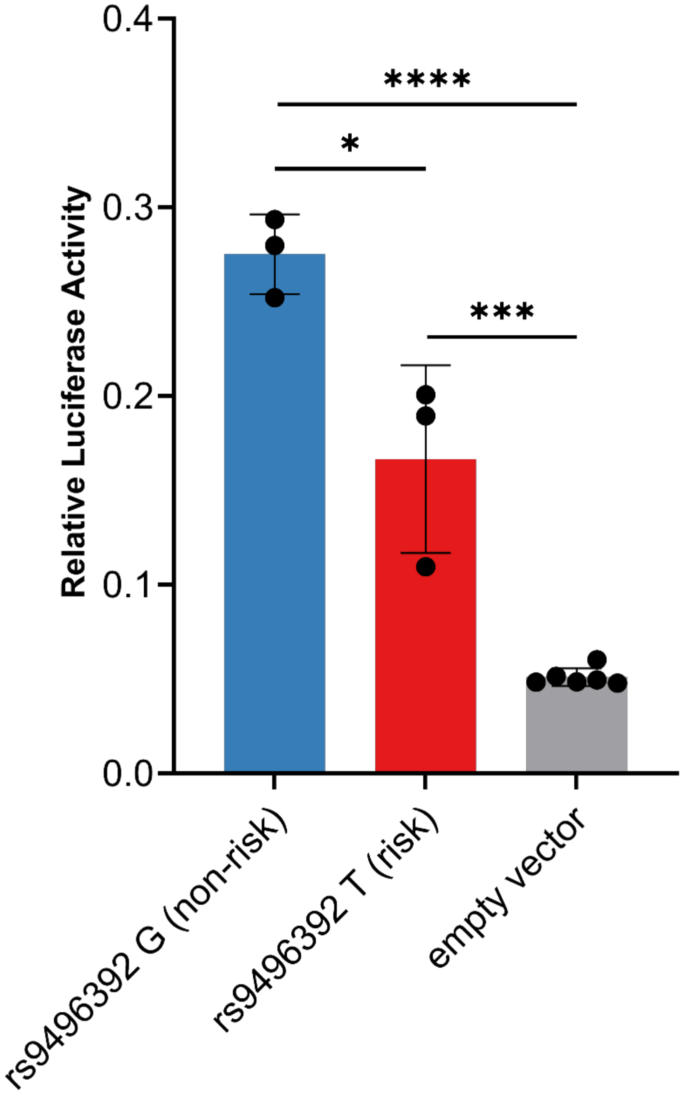


**Supplementary Figure 8.** Luciferase assay results of three additional biological replicates comparing allelic enhancer activity of rs9496392. *, $p < 0.05$; ***, $p < 0.001$; ****, $p < 0.0001$.

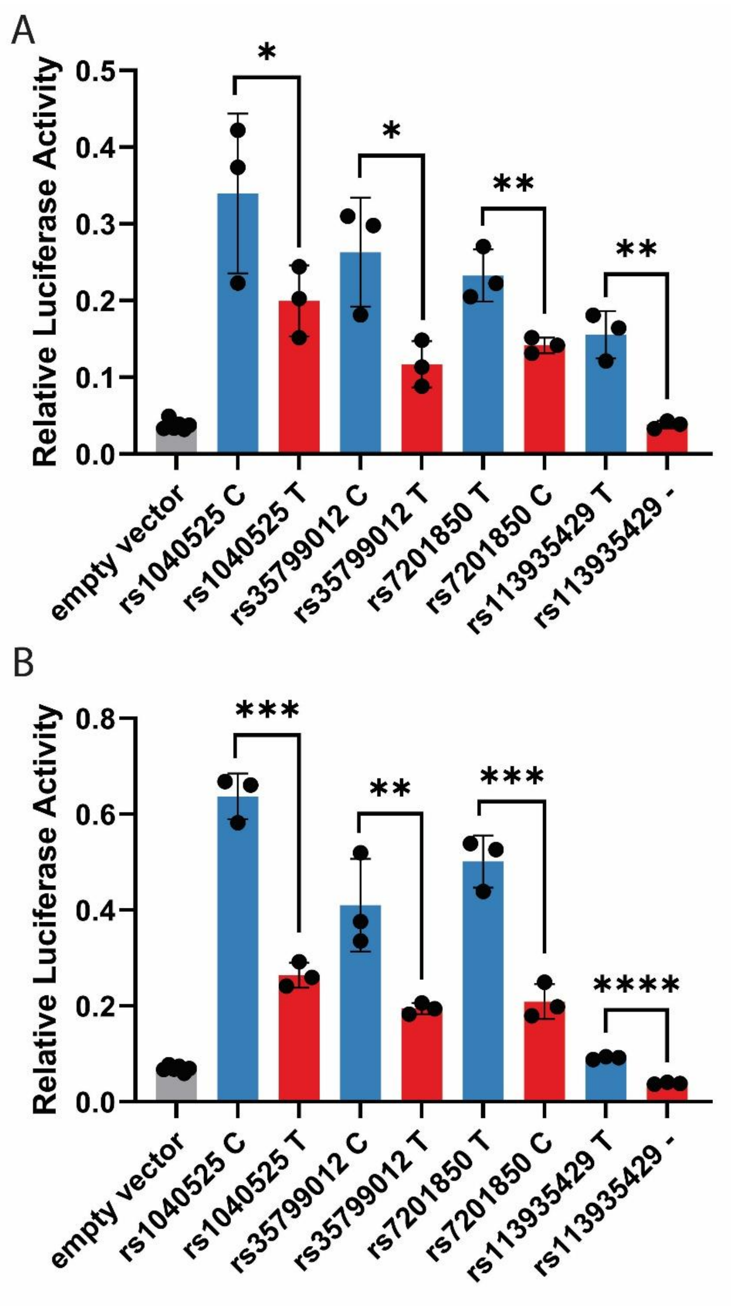


**Supplemental Figure 9.** Allele-specific relative luciferase activity of rs1040525, rs35799012, rs7201850, and rs113935429 in TC28a2 cells. A) Three biological replicates confirming differential activity. B) Three additional, independent biological replicates to confirm differential activity. Blue columns indicate reference alleles and red columns indicate alternate alleles. *, $p < 0.05$; **, $p < 0.01$; ***, $p < 0.001$. Statistical significance between CRS activity and empty-vector negative controls was determined by one-way ANOVA; statistical significance between SNP alleles was determined by unpaired Student's *t*-test.

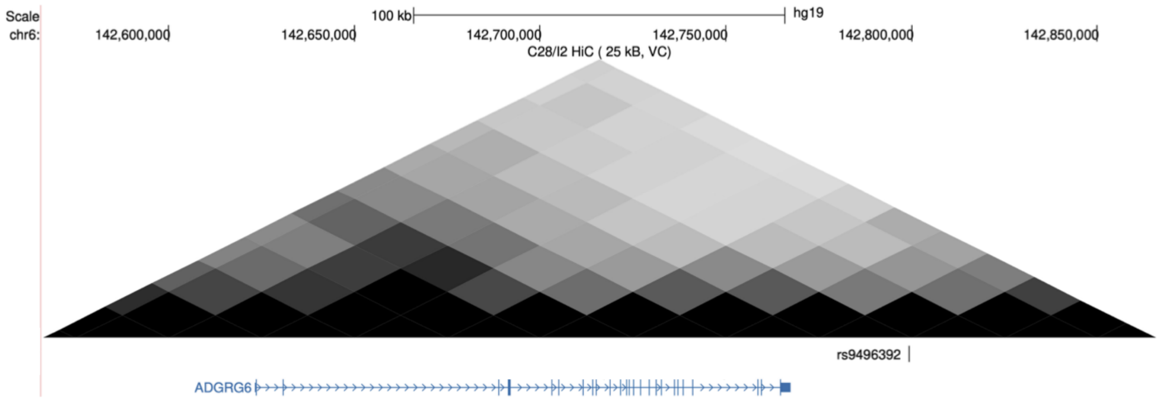


**Supplemental Figure 10.** rs9496392 3D genome interactions throughout the *ADGRG6* locus in C28/I2 chondrocytes.
